# Not all seabird species can overcome marine threats when predator removal at their colonies is prioritised

**DOI:** 10.1101/770263

**Authors:** Stephanie B. Borrelle, Holly P. Jones, Yvan Richard, Roberto Salguero-Gómez

## Abstract

Seabirds are globally threatened. In the face of multiple threats, it is critical to understand how conservation strategies that mitigate one threat intersect with others to impact population viability. Marine threats, including pollution, climate change, and fisheries could derail gains to seabird populations resulting from arduous predator eradication efforts. However, this potentially negative effect is yet to be evaluated. We test whether mortality from marine threats can subvert the on-going recovery of 17 seabird species from 37 colonies on islands worldwide where predators were removed. We use demographic modelling to estimate potential adult mortality from fisheries, plastic ingestion, and climate change. For 82% of the species we examine, marine threats do not impede recovery following predator eradication. However, for six colonies of three species, *Calonectris diomedea, C. borealis*, and *Ardenna carneipes*, mortality from multiple marine threats may interrupt their recovery. Combining our demographic approach with comparative phylogenetic methods, we explore whether foraging niche, range, and morphometric traits inform the vulnerability to marine threats using an expanded dataset of 81 seabird species. Our analyses reveal surface filtering and pursuit diving species, and species with smaller at-sea distributions to be most vulnerable to declines due to multiple threats. However, these traits do not necessarily predict species’ vulnerability to marine threats in the absence of predators at nesting colonies, suggesting that shared traits may not be useful to infer vulnerability to multiple marine threats. Post-eradication monitoring to determine whether species require additional conservation management following predator eradication are essential in the face of intensifying pressures in the marine environment.

---

Seabirds are one of the most globally threatened taxa due to a plethora of threats, including habitat alteration, invasive species, and pollution (IUCN, 2017; Rodríguez et al., 2019). These threats can act simultaneously (Barbraud et al., 2012; Lawler et al., 2002), be separated by time and/or space (Sutherland et al., 2012), or target different stages in a species’ life cycle (Votier, Hatchwell, Mears, & Birkhead, 2009). Pelagic seabirds’ (order Procellariiforme; *i.e.*, albatrosses, petrels, shearwaters, prions, and fulmars; hereafter ‘seabirds’) typically high adult survival and low intrinsic population growth rates render their populations vulnerable to anthropogenic stressors because even small decreases in adult survival can potentially affect their long-term reproductive output (Schreiber & Burger, 2002). While seabirds exhibit variation in life-history traits (*e.g.*, longevity, age at maturation, reproductive success), trophic relations, and geographic distributions, these species tend to be extremely vulnerable to (i) invasive mammalian predators at their breeding sites (Jones et al., 2008; Towns et al., 2011), and (ii) anthropogenic marine stressors, such as pollution, fisheries, and climate change (Provencher et al., 2018; Rodríguez et al., 2019).

To date, over 1,000 islands worldwide have been rid of invasive predators by management actions, leading to significant seabird population gains (Brooke et al., 2018; Jones et al., 2016). But are predators on islands seabirds’ main cause of vulnerability? Marine threats, such as fisheries bycatch, plastic pollution, and unpredictable impacts of climate change can contribute to mortality. These agents can cause severe seabird population reductions (Genovart et al., 2017), even in the face of positive responses from predator eradication. Quantifying the population-level impacts of single or multiple marine threats can be challenging because we lack detailed demographic information for most species (Richard, Abraham, & Filippi, 2017; Rodríguez et al., 2019). Indeed, monitoring following predator eradication is sparse or absent, belying an inability to accurately assess species recovery rates (Brooke et al., 2018; Buxton, Jones, Moller, & Towns, 2014; Kappes & Jones, 2014). This uncertainty is confounded by (i) the challenges of quantifying at-sea mortality, (ii) sub-lethal effects of marine stressors on population viability (e.g., plastic ingestion Clukey et al., 2018; Lavers, Bond, & Hutton, 2014; Oro, 2014; Tanaka et al., 2015), and (iii) the lack of understanding regarding how multiple stressors interact or cumulate to affect seabird population trends (Burthe, Wanless, Newell, Butler, & Daunt, 2014). Such limitations make it challenging to address how current conservation strategies that mitigate one threat intersect with other threats to impact seabird population viability.

Here, we explore how multiple marine threats may impact the recovery of seabird populations that have experienced predator eradication at their colonies, and ask whether seabirds’ phylogenetically conserved traits can help predict their vulnerability to marine and terrestrial threats. We hypothesise that (1) high adult mortality caused by fisheries bycatch, plastic ingestion, and climate change (marine threats, hereafter) impedes the recovery of seabirds following predator eradication, and that (2) the vulnerability of a species to marine threats can be inferred from phylogenetically shared morphometric and ecological traits. The latter hypothesis, if supported, would provide an approach to infer risk from closely related species where adult mortality data from marine threats is lacking.

## Methods

We estimate species vulnerability by calculating the annual mortality threshold, that is, the number of adults in a population that can be killed annually with the population still remaining viable. We then calculate the risk of local population extinction to each of the aforementioned marine threats of 17 seabird species from 37 colonies on 24 islands around the world where predators have been eradicated (Table 1, inset map in Figure 1).

**Table 1.** For 82% (6 of 37) of the colonies of seabirds we evaluated, mortality from marine threats do impede recovery of the natural populations following predator eradication^1^.

| Species (IUCN status) | Colony (region) | Colony Population size<br>(sd) | Annual mortality<br>threshold (sd) | Fisheries<br>mortality<br>(sd) | Plastic<br>mortality<br>(sd) | Climate<br>mortality<br>(sd) | Risk ratio<br>(sd) |
| --- | --- | --- | --- | --- | --- | --- | --- |
| *Calonectris borealis* (LC) | La Scola (Mediterranean) | 131 (165) | 5 (5.8) | 3.28 (4.61) | 0.12 (0.02) | 0.14 (0.02) | 3.55 (4.62) |
| *Calonectris diomedea* (LC) | Sand Island (Atlantic) | 4,316 (5,953) | 215 (314.9) | 2.68 (4.61) | 0.09 (0.02) | 0.11 (0.02) | 2.88 (4.62) |
| *Ardenna carneipes* (NT) | Lady Alice (New Zealand) | 3,510 (5,174) | 118 (176.0) | 0.98 (1.49) | 0.12 (0.02) | 0.15 (0.02) | 1.25 (1.50) |
|  | Ohinau (New Zealand) | 7,367 (10,459) | 246 (342.1) | 0.94 (1.38) | 0.12 (0.02) | 0.15 (0.02) | 1.22 (1.38) |
|  | Whatupuke (New Zealand) | 4,196 (5,991) | 139 (194.8) | 0.91 (1.26) | 0.12 (0.02) | 0.15 (0.02) | 1.19 (1.26) |
|  | Coppermine (New Zealand) | 4,723 (6,421) | 185 (258.8) | 0.81 (1.18) | 0.10 (0.01) | 0.13 (0.01) | 1.04 (1.18) |
| *Ardenna pacifica* (LC) | Oahu (Hawaii) | 5,811 (9,688) | 256 (438.8) | 0.74 (1.14) | 0.05 (0.01) | 0.12 (0.02) | 0.91 (1.14) |
|  | Kadomo (Pacific) | 1,377 (2,0212) | 61 (91.3) | 0.73 (1.03) | 0.05 (0.01) | 0.12 (0.02) | 0.90 (1.04) |
|  | Serrurier (Australia) | 60,806 (85,885) | 2681 (3943.7) | 0.71 (1.05) | 0.05 (0.01) | 0.12 (0.02) | 0.88 (1.06) |
|  | Surprise (SW Pacific) | 511 (689) | 22 (31.4) | 0.70 (1.08) | 0.05 (0.01) | 0.12 (0.02) | 0.87 (1.08) |
|  | Monuriki (Pacific) | 3,505 (4,811) | 154 (220.2) | 0.69 (0.91) | 0.05 (0.01) | 0.12 (0.02) | 0.86 (0.91) |
|  | Jarvis (Central Pacific) | 167 (223) | 7 (10.2) | 0.69 (0.87) | 0.05 (0.01) | 0.12 (0.02) | 0.86 (0.88) |
| *Puffinus puffinus* (LC) | Lundy (UK) | 500 (639) | 11 (16.5) | 0.00 (0.00) | 0.21 (0.11) | 0.27 (0.14) | 0.49 (0.25) |
|  | Ramsey (UK) | 1,416 (2,155) | 33 (56.7) | 0.00 (0.00) | 0.21 (0.10) | 0.27 (0.13) | 0.48 (0.24) |
| *Pterodroma hypoleuca* (LC) | Selvagem Grande (North Atlantic) | 1,743,481 (2,401,203) | 45177 (67917.4) | 0.00 (0.00) | 0.19 (0.05) | 0.20 (0.05) | 0.39 (0.09) |
| *Pterodroma pycrofti* (VU) | Red mercury (New Zealand) | 35,222 (54,520) | 912 (1443.2) | 0.00 (0.01) | 0.17 (0.04) | 0.20 (0.05) | 0.37 (0.09) |
|  | Aorangi (New Zealand) | 133 (178) | 3 (4.6) | 0.00 (0.01) | 0.16 (0.04) | 0.20 (0.05) | 0.37 (0.09) |
| *Pterodroma cookii* (VU) | Hauturu (New Zealand) | 43,165 (58,440) | 1095 (1527.3) | 0.00 (0.00) | 0.16 (0.02) | 0.20 (0.03) | 0.36 (0.05) |
| *Pterodroma gouldi* (LC) | Hauturu (New Zealand) | 98 (125) | 3 (3.5) | 0.01 (0.01) | 0.12 (0.03) | 0.20 (0.05) | 0.33 (0.08) |
|  | Burgess (New Zealand) | 2562 (3561) | 66 (94.5) | 0.01 (0.02) | 0.11 (0.03) | 0.20 (0.05) | 0.33 (0.08) |
|  | Atihau (New Zealand) | 10,280 (14,348) | 266 (358.3) | 0.01 (0.02) | 0.11 (0.03) | 0.20 (0.05) | 0.33 (0.08) |
|  | Motuharekeke (New Zealand) | 2,571 (3,608) | 66 (93.3) | 0.01 (0.01) | 0.11 (0.03) | 0.20 (0.05) | 0.33 (0.08) |
|  | Otata (New Zealand) | 814 (1045) | 21 (26.7) | 0.01 (0.02) | 0.11 (0.03) | 0.20 (0.05) | 0.33 (0.08) |
|  | Moutohora (New Zealand) | 69,695 (96,614) | 1812 (2641.2) | 0.01 (0.02) | 0.11 (0.03) | 0.20 (0.05) | 0.33 (0.08) |
| *Puffinus assimilis* (LC) | Burgess (New Zealand) | 1,143 (1,775) | 47 (73.3) | 0.00 (0.00) | 0.12 (0.02) | 0.13 (0.02) | 0.24 (0.03) |
|  | Red mercury (New Zealand) | 3,350 (4,524) | 137 (184.7) | 0.00 (0.00) | 0.12 (0.02) | 0.13 (0.02) | 0.24 (0.03) |
|  | Atihau (New Zealand) | 1,161 (1,607) | 48 (67.9) | 0.00 (0.00) | 0.12 (0.02) | 0.12 (0.02) | 0.24 (0.03) |
| *Ardenna bulleri* (VU) | Aorangi (New Zealand) | 25,261 (33,957) | 872 (1201.5) | 0.00 (0.00) | 0.09 (0.01) | 0.15 (0.02) | 0.24 (0.03) |
| *Puffinus gavia* (LC) | Atihau (New Zealand) | 1,055 (1,497) | 43 (60.5) | 0.01 (0.01) | 0.11 (0.02) | 0.13 (0.02) | 0.24 (0.03) |
|  | Hauturu (New Zealand) | 253 (345) | 10 (14.5) | 0.01 (0.01) | 0.10 (0.01) | 0.13 (0.02) | 0.23 (0.03) |
| *Pterodroma ultima* (NT) | Oeno (S Pacific) | 22,037 (28,992) | 957 (1271.6) | 0.01 (0.01) | 0.09 (0.01) | 0.12 (0.01) | 0.21 (0.03) |
| *Ardenna grisea* (NT) | Lady Alice (New Zealand) | 95 (117) | 5 (5.7) | 0.00 (0.00) | 0.09 (0.01) | 0.11 (0.01) | 0.20 (0.02) |
| *Fregetta maoriana* (CR) | Hauturu (New Zealand) | 7,906 (11,148) | 559 (798.7) | 0.01 (0.02) | 0.05 (0.01) | 0.07 (0.01) | 0.14 (0.03) |
| *Pelecanoides urinatrix* (LC) | Atihau (New Zealand) | 2,524 (3,141) | 205 (256.6) | 0.00 (0.01) | 0.03 (0.00) | 0.06 (0.01) | 0.09 (0.01) |
|  | Burgess (New Zealand) | 70,153 (95,301) | 5693 (7819.0) | 0.00 (0.01) | 0.03 (0.00) | 0.06 (0.01) | 0.09 (0.01) |
|  | Hauturu (New Zealand) | 814 (1,004) | 66 (83.2) | 0.00 (0.01) | 0.03 (0.00) | 0.06 (0.01) | 0.09 (0.01) |
| *Pelagodroma marina* (LC) | Burgess (New Zealand) | 43,617 (58,281) | 3388 (4562.2) | 0.00 (0.00) | 0.01 (0.00) | 0.07 (0.01) | 0.08 (0.01) |

<sup>1</sup>
<sup>1</sup> Model estimates of population size, the annual mortality threshold, and the potential mortality from each of the marine threats of fisheries, climate change, and plastic ingestion is given as the mean and standard deviation of the mean. The high risk (>1) for *Calonectris borealis*, *C. diomedea*, and *Ardenna carneipes* was driven predominantly by mortality from fisheries. The risk ratio was calculated as potential mortalities yr<sup>-1</sup> / annual mortality threshold (Richard & Abraham, 2013); when this risk ratio ≥1, adult mortality from each of the evaluated threats may impede the recovery of a colony even after predator eradication. For the remaining 37 colonies of 17 species that we analysed, the risk ratio was <1, implying that marine threats are not impeding the recovery of these populations following predator eradication.

**Fig. 1.**
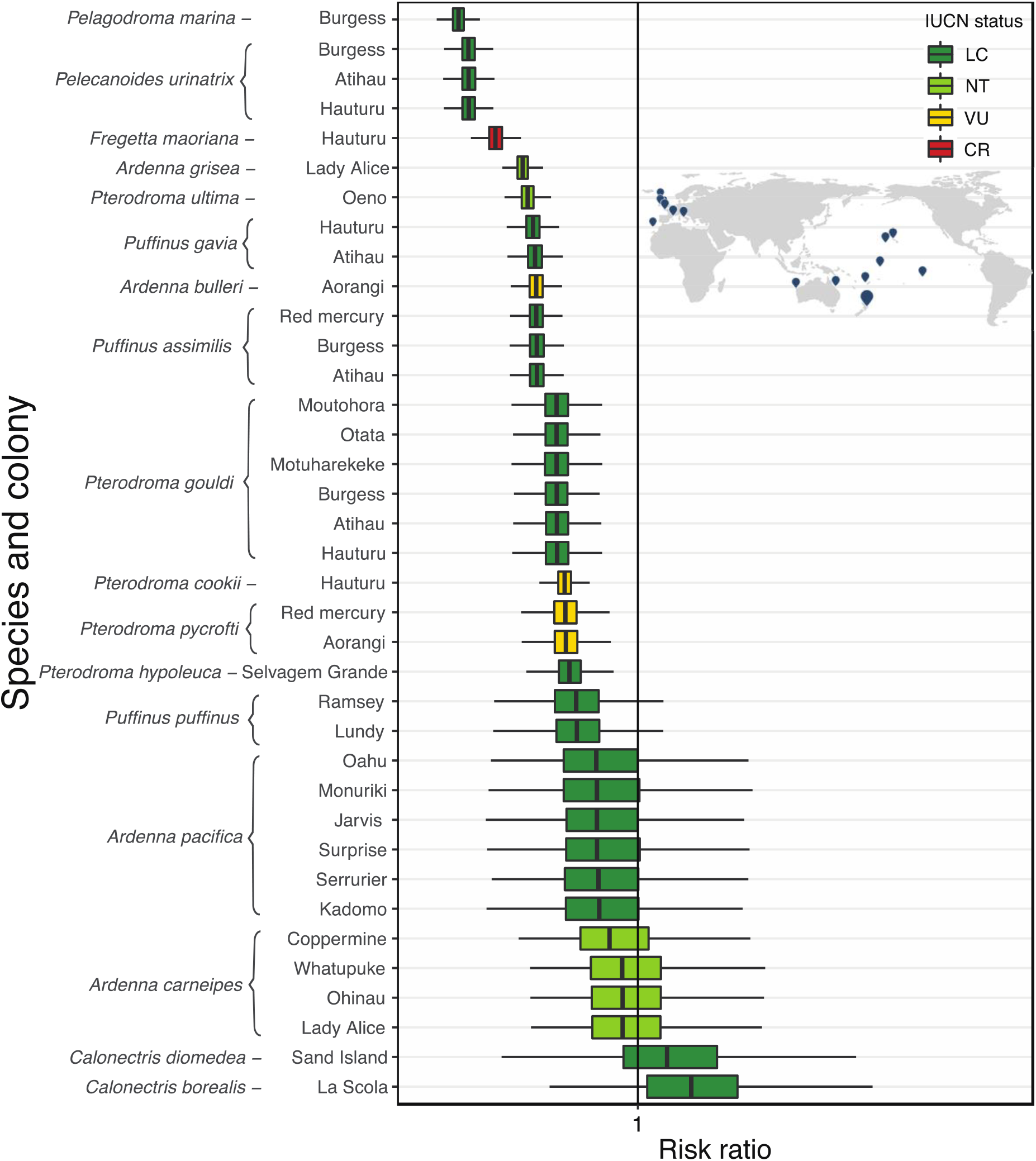
The risk ratio for the 37 colonies of 17 species (map inset shows colony locations worldwide), on 24 islands where invasive predators have been eradicated. The risk ratio was calculated as potential mortalities yr^−1^/ annual mortality threshold (Richard & Abraham, 2013); when this risk ratio ≥1, adult mortality from each of the evaluated threats may impede the recovery of a colony even after predator eradication. Colours for each species correspond to the IUCN Red List status: LC Least Concern in dark green; NT Near threatened in light green; VU Vulnerable in yellow; CR Critically Endangered in red.

To calculate the annual mortality threshold and the risk ratio, we used the Demographic Invariant Method (DIM, hereafter). The DIM requires minimal demographic information to estimate (i) the intrinsic annual population growth rate of a species under optimal conditions, and (ii) the annual mortality threshold for a given population. The annual mortality threshold corresponds to the maximum number of individuals in a population that could be *extracted* annually, *e.g.*, due to any source of mortality. Using the DIM approach, we estimate the annual mortality threshold for 81 seabird species (order: Procellariiformes) with equation 1 (*20*)(Richard et al., 2017):

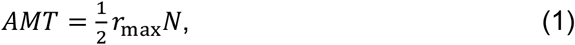

where *r*_max_, or the population growth rate above replacement per generation is (Niel & Lebreton, 2005; Richard et al., 2017) calculated from equation 2:

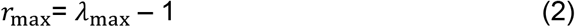

The Demographic Invariant Method (DIM) combines matrix population models and allometric relationships to calculate the intrinsic annual population growth rate, *λ*_max_ or maximum annual growth rate (Dillingham et al., 2016; Niel & Lebreton, 2005; Richard et al., 2017). The model estimates of mean age at maturation (α), adult survival (*s*), and population size *(N*) used to calculate *λ*_max_ and the annual mortality threshold were defined by the mean and standard deviation from data obtained from the BIDDABA database (Lebreton & Gaillard, 2017; unpublished) and the COMADRE Animal Matrix Database (Salguero Gómez et al., 2016). We also used the peer-reviewed literature and online data sources (Birdlife International: birdlife.org/datazone/species and New Zealand Birds Online: nzbirdsonline.org.nz; Online Material; Table S1).

For age at maturation *α*, samples were drawn from the gamma distribution with the shape and rate calculated using the function *gamma.parms.from.quantiles* (Joseph & Bélisle, 2012), and the function *rgamma* from the *base R* package (R Core Team, 2013). For adult survival *s*, the mean and standard error were derived by calculating the logit of the mean, and then back transformed. The standard deviation of the logit of the mean for *s*, was calculated using the Delta method, that is a general method to derive the variance of a function, shown in equation 3 (Richard & Abraham, 2013):

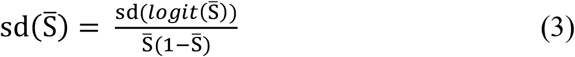

The required demographic information to calculate *λ*_max_ are age at first reproduction (*α*), adult survival (*s*), and an allometric constant *a*_rT_ = 1, *sensu* (Niel & Lebreton, 2005), with equation 2:

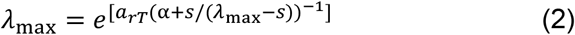

Population data are needed to calculate the annual mortality threshold; however, because census data for seabirds are typically from colony surveys where only breeding adults can be counted, the total population size represented by the census data is often unknown (Dillingham et al., 2016; Richard & Abraham, 2013). Further, immature birds spending their first few years at sea before returning to the colony to breed, and some adults taking sabbaticals from breeding make this accurate calculation more challenging (Richard & Abraham, 2013; Warham, 1990). We estimated the mean and standard deviation of the population size assuming a normal distribution in R (R Core Team, 2013). We set a standard deviation to 0.05 to account for the uncertainty in population estimate data (see Online Material for further discussion on data quality). Data were log-transformed to meet statistical assumptions.

### Marine threat impact

The first goal in our study is to examine the impacts of marine stressors on population recovery for 37 colonies including 17 species on 24 islands where predators have been eradicated, where we were able to find demographic, and threat impact data (Online material; Figure S1). By using this subset, we assume that the populations were minimally affected by terrestrial threats (i.e., recovering from the pressure of invasive mammalian predators following predator removal, minimal density-dependence effects, no limit of available habitat, and no resource limitations) (Borrelle, Boersch-Supan, Gaskin, & Towns, 2016; Ismar et al., 2014; Niel & Lebreton, 2005). The following details the estimation of annual mortality for the 17 species by three key agents in our model:

#### Fisheries by-catch

We used the mean annual potential mortality from fisheries from Richard et al (Richard et al., 2017) for 12 of the seabird species included in the colony analysis (Appendix 2). The estimates of annual potential fatalities come from the New Zealand Ministry for Primary Industries bycatch data of four fishing methods; trawl, surface and bottom long-line, and set nets from the New Zealand exclusive economic zone (EEZ) for the period 2006-07 to 2014-15. Estimates for *Calonectris diomedea* were from Belda & Sanchez (Belda & Sanchez, 2001) and we used the same estimate for the closely related species C. borealis, which may underestimate the impact of fisheries bycatch for this species (Online material). The species *Pterodroma hypoleuca*, *P. ultima* and *Puffinus puffinus* were assumed to be low risk from fisheries because are not highly reported as bycatch in the literature (IUCN, 2017). We assumed that the proportion of adults in the total population that will potentially be killed by fisheries would be the same proportion for each colony we evaluated. To account for the uncertainty in these estimates, we sampled the mean and 95% credible intervals from a log-normal distribution using 5,000 bootstraps (Table 2).

#### Plastic Pollution

We searched Google Scholar and Web of Science using the terms seabird species + plastic ingestion (e.g. “*Ardenna carneipes*” and “*plastic* and ingestion*”). We calculated the proportion of the population at risk from mortality due to ingesting plastics by the average frequency of occurrence of plastic reported in plastic ingestion studies for each species (Avery-Gomme et al, *In review*; Online materials; Appendix 1). We assumed that 0.5% of individuals died as a result of ingesting plastics, we sampled the mean and 95% credible intervals from a log normal distribution using 5,000 bootstraps (Online material; Table S2). To account for the uncertainty in the estimate of the proportion of individuals killed from ingesting plastic, we tested the sensitivity to 1% and 5% mortality (Online material; Figs. S3 & S4).

#### Climate change

We assume that the annual mortality rate due to the direct impacts of climate change was 0.5% for all populations of the 17 studied species, in the absence of peer-reviewed estimates. In order to account for the high uncertainty in this estimate a sample of 5,000 bootstraps of the mortality rate due to climate change was drawn from a log-normal distribution of mean 0.005 and standard deviation of 0.05 (Online material: Table S2). To test the sensitivity of our results to the assumed value, the calculations were repeated with a mean of 0.01 and 0.05 (Online material; Figs. S5 & S4). The impacts on climate change, should therefore be considered as a scenario model rather than empirical assessment (Online Material; Table S3). See Online Material for further discussion on uncertainty.

To calculate the proportional per capita mortality rate for each of the colonies evaluated, we used the annual potential mortalities from each of the threats / total population. We then calculated the risk of a species to each of the above stressors and combined potential mortality from all threats together. The risk ratio is calculated as potential mortalities yr^−1^/ annual mortality threshold (Richard & Abraham, 2013). A ratio close to one or above means that the species is at high risk of ‘over-harvesting’ by marine stressors. All statistical analyses were carried out in *R* (R Core Team, 2013).

##### Predictors of vulnerability

The second goal of our study was to test whether factors informing the ecological niche of a species inform its vulnerability to marine threats. To do this we use phylogenetic generalised least squares (PGLS) regression to test whether the key morphometric traits (body size) and ecological seabird characteristics (foraging strategy, diet, and at-sea distribution) of the studied species can be used to predict vulnerability, using the annual mortality threshold as a proxy. Due to shared ancestry, closely related species are expected to share similar trait values (Symonds & Blomberg, 2014). To quantify the phylogenetic signal of our traits of interest: body mass, foraging strategy, diet, and at-sea distribution, we estimated Pagel’s *λ* (not to be confused with the intrinsic population growth rate *λ*_max_), a scaling parameter for the phylogenetic correlation between species that ranges from 0 (no role of phylogeny in determining trait variation) to 1 (trait variation fully explained by phylogeny assuming Brownian motion (Freckleton, Harvey, & Pagel, 2002). We used an expanded dataset that included 81 pelagic seabird species, where demographic parameters were available to calculate the annual mortality threshold using the methods described above. Next, we obtained the bird phylogeny by Jetz et al. (2012), which contains time-calibrated phylogenetic relationships from conserved regions of the genomes of 9,993 extant bird species. We manipulated the tree to prune it to our expanded dataset of 81 seabird species to calculate Pagel’s *λ* using the *R* packages *phytools* (Revell, 2012)*, ape* (Paradis, Claude, & Strimmer, 2004), and *caper* (Orme, 2013).

To examine the ecological and morphometric metrics that best predict the risk of a species to anthropogenic sources of at-sea mortality, we used phylogenetic generalised least squares (PGLS) regression. We used the annual mortality threshold as our response variable and our set of explanatory variables to morphometric (mean adult body size), and ecological niche (foraging strategy, prey type, at-sea distribution). Since evidence exists that these are indicators of species’ likelihood to interact with fishing vessels (Genovart et al., 2017), ingest plastic pollution (Day, Wehle, & Coleman, 1985; van Franeker & Law, 2015), and/or their overall risk to extinction (and spatial distribution 30, 33, e.g., body size 37). The PGLS were done with the function *pgls* of the *caper R* package (Orme, 2013), and using the bird phylogeny described above as the backbone.

## Discussion

Marine threats do not affect the population recovery of 82% (14 out of 17) of the species in our risk analysis. However, for six colonies including three closely related species -*Calonectris diomedea* (Cory’s Shearwater)*, C. borealis* (Scopoli’s shearwater), and *Ardenna carneipes* (Flesh-footed shearwater), marine threats are an important risk to post-eradication recovery (Table 1). Mortality from fisheries is the predominant threat driving this pattern for all of these species (Figure 2, Table 1). Indeed, all of these species are often reported as bycatch in recreational and commercial fishing activities (Belda & Sanchez, 2001; Genovart et al., 2017; Richard et al., 2017), but they may be highly vulnerable to plastic ingestion (Lavers et al., 2014). Such impacts are compounded by potential climate-induced changes to prey distributions, leading to the potential need to travel longer distances to forage than previously recorded, e.g., (Bost et al., 2015).

**Fig. 2.**
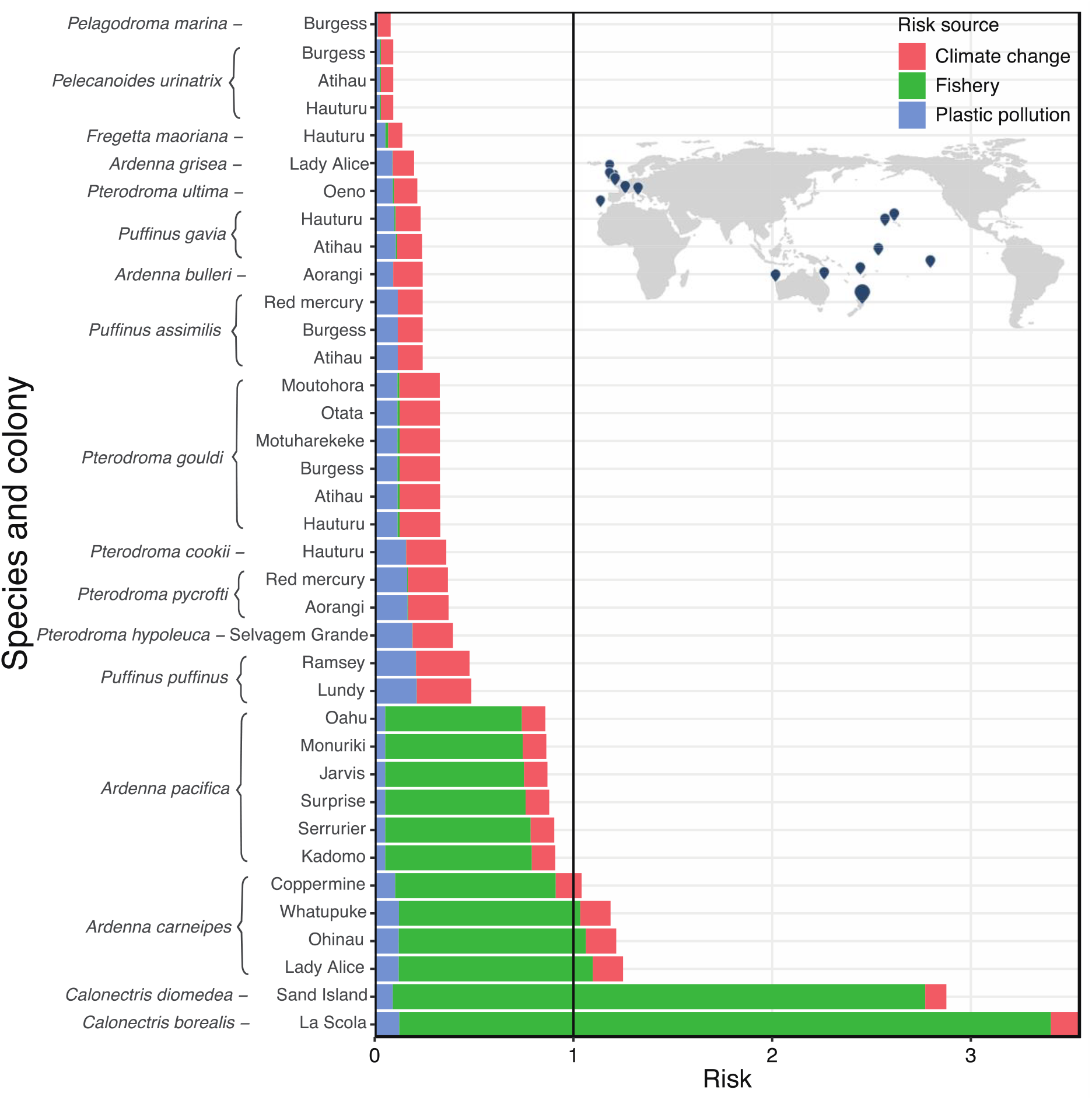
The relative contributions of the marine threats of mortality from climate change, fisheries bycatch, and plastic ingestion for the colonies (*n=37*, map inset show location) of 17 species, on 24 islands where predators have been eradicated. The risk ratio was calculated as potential mortalities yr^−1^/ annual mortality threshold (Richard & Abraham, 2013); when this risk ratio ≥1, adult mortality from each of the evaluated threats may impede the recovery of a colony even after predator eradication.

Wedge-tailed shearwater’s (*Ardenna pacifica*) are also at relatively high risk to marine threats, despite having a risk ratio lower than 1 (mean=0.88 ± 0.01; Table 1). Indeed, if plastic pollution or climate change result in higher mortality than we have estimated here (Online materials; Fig. S3 & S4), declines would likely continue for *A. pacifica*, even after predator eradication (IUCN, 2017). In fact, any of the populations we analysed were to undergo an average of 5% mortality from marine threats, recovery for all but three of the smallest species we analysed would be impeded (Online material; Fig. S4 & S6). Thus, it is crucial to understand how multiple marine threats impact seabird populations following predator eradication. Such information will help to avoid declines as these pressures are likely to intensify.

### Inferring vulnerability to multiple threats

With an expanded dataset including 81 seabirds, we use phylogenetic comparative analyses to explore whether key morphometric (body size) and ecological characteristics (primary foraging strategy, primary diet, and at-sea distribution) predict species’ vulnerability to declines, using the annual mortality threshold as a proxy. We validate our metrics of population performance and vulnerability against the IUCN Red List categories of those species (Figure 3.A). The annual mortality threshold estimates for the 81 species in our analysis are congruent with IUCN Red List threat categories. Figure 3.A shows a negative correlation between threat status of a species and its annual mortality threshold. Our phylogenetic analyses examined species traits that might predict the annual mortality threshold of the 81 species, and retained at-sea distribution and biomass (Figure 3.B), and the foraging strategies of surface filtering and pursuit diving (online materials; Table S3). However, the same traits did not fully inform risk when compared with colony-level risk analysis data (Table 1). For example, based on the phylogenetic analysis, we would predict that *Pterodroma gouldi* (Grey-faced petrel), which is a surface forager, has a small distribution, and large body size, to have a high risk ratio (Table 1; online materials; appendix 1). However, the risk ratio for all six of the *P. gouldi*’s colonies are 0.33 (sd: 0.08; Table 1). Further, research shows *P. gouldi* population increases where predators have been removed or are controlled (Buxton et al., 2014; Greene, Taylor, & Earl, 2015). In addition, this species is not reported to be at high risk from fisheries bycatch or plastic ingestion (Richard et al., 2017).

**Fig. 3.**
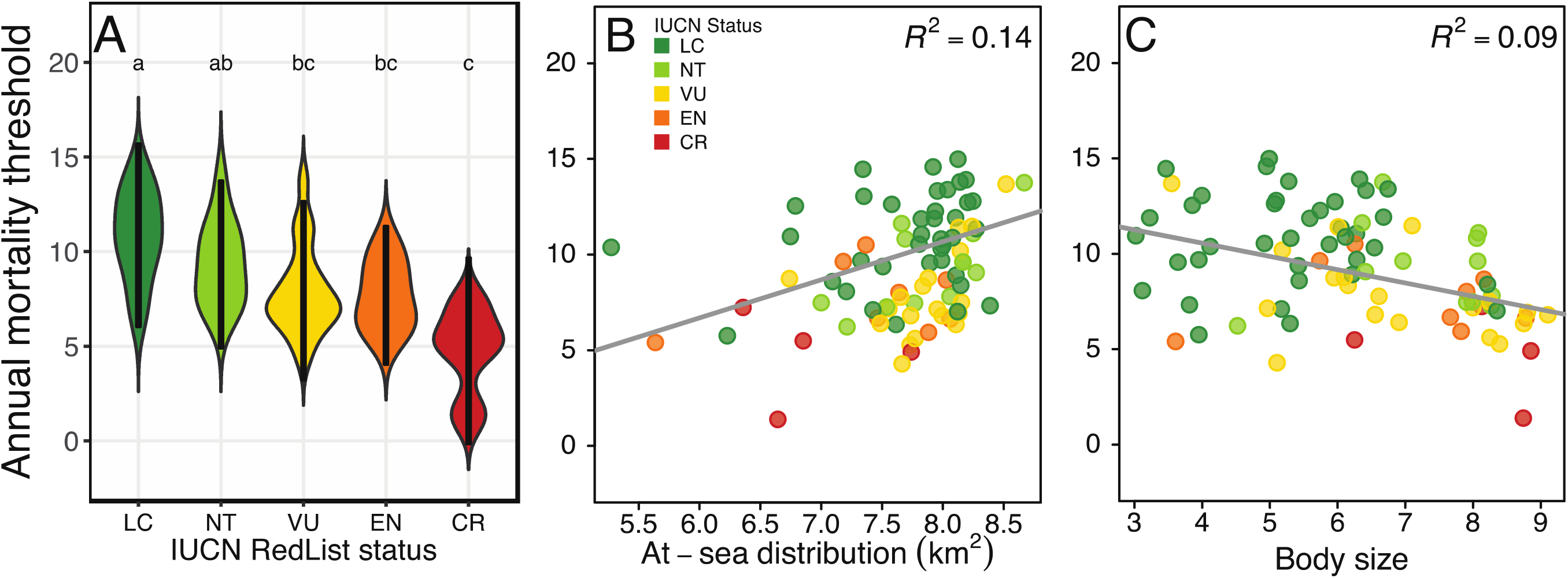
The annual mortality threshold serves as a basis to (A) predict the IUCN RedList status of the examined 81 seabird species, and it is also predicted by key ecological predictors such as (B) seabird species at-sea distribution (C) and adult body mass. The annual mortality threshold is the maximum number of breeding adults that can be removed annually from a population without causing it to decline. Groupings correspond to the IUCN Red List status: LC Least Concern in dark green; NT Near threatened in light green; VU Vulnerable in yellow; EN Endangered in orange; CR Critically Endangered in red. Letters on top of each IUCN group in panel A are post-hoc Tukey scores; when two groups do not share the same letter, their annual mortality threshold scores are statistically different at *P*<0.05 (supporting material; Table S2). (B) and (C) contain the phylogenetic generalised least squares for the relationship between the annual mortality threshold and at-sea distribution (log-scale); Pagel’s λ=0.83, 95% CI 0.42-0.98, *F-ratio=*19.14, df=79, *P*<0.001), and adult body mass (Pagel’s *λ*=0.52, 0.06-0.87, *F-statistic=*4.05, df=79, *P*<0.001).

Surface filtering and pursuit diving seabirds are more vulnerable to declines than species with other foraging strategies (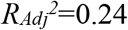; *f*=13.18; *P*_*Adj*_<0.001; and 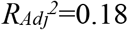, *f*=11.18; *P*_*Adj*_<0.001, respectively; online materials; Table S3). This relationship is highly preserved among closely related species (Pagel’s *λ*=1.00, 0.44-1.00; Pagel’s *λ*=1.00, 0.64-1.00, respectively). Surface foraging species have been identified as at high risk for ingesting plastic in our phylogenetic analysis, and by other authors (Roman, Bell, Wilcox, Hardesty, & Hindell, 2019) (Table 1). All the species in our colony-level analysis are surface foragers or pursuit divers (online materials; Appendix 1), yet only three species are vulnerable to marine threats in the absence of invasive predators at their breeding sites. Foraging strategy and prey type have been previously linked to seabirds’ propensity to ingest plastic (Day et al., 1985; Provencher, Gaston, Mallory, O’hara, & Gilchrist, 2010; Ryan, 1987). Longline and gillnet fisheries are dominant sources of adult mortality for surface filtering and pursuit diving species (Bærum et al., 2019; Rodríguez et al., 2019). Relating specific foraging strategy to climate change risk is challenging; however, species that are flexible in foraging strategy, allowing shifts in prey with changing conditions may be more resilient to the impacts of climate change (Riou et al., 2011). While foraging strategy certainly plays a role, the risk of adult mortality from marine threats is also likely to be strongly influenced by spatial co-occurrence, individual behaviour (Krüger, Pereira, Paiva, & Ramos, 2019), and distribution of both stressors and birds (Ryan, 1987, 2016).

Species with smaller at-sea distributions had lower annual mortality thresholds than those with larger distributions (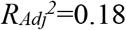; *t*=3.7; *P*<0.001; Figure 3.B). The vulnerability of a species and their spatial distribution is conserved among closely related species (Pagel’s λ=0.83, 0.42-0.97; online materials; Table S3). Our finding that the vulnerability of a species is influenced by distribution is in agreement with the inclusion of range size as one of the main metrics of IUCN Red List evaluations of threatened status (IUCN, 2017). However, for the species with the highest risk to marine threats following predator eradication, the opposite is true – species with large at sea distributions are more at risk.

Theory predicts that species with large range sizes may be more resilient to environmental stressors and adult mortality (Cooke, Eigenbrod, & Bates, 2019; Paniw, Ozgul, & Salguero Gómez, 2018; Sæther & Engen, 2010). Large at-sea distributions mean that when environmental or foraging conditions are poor, highly mobile species, such as seabirds, may be able to move to more favourable foraging or breeding grounds. During these unfavourable periods, adult survival can remain high, but reproductive success may be low (Giudici, Navarro, Juste, & González-Solís, 2010; Weimerskirch, 2001). However, three closely related species in our analyses, all with large at-sea ranges, have the highest risk for marine threats following predator eradication. The Atlantic colony of *C. diomedea*, the Mediterranean colony of *C. borealis*, and all four of the New Zealand colonies of *A. carneipes*, have range sizes of over 74, >74.3, and 180 million km^2^, respectively (Birdlife International, 2016; Jetz et al., 2012) (Figure 1; Table 1). Indeed, other research has found that Mediterranean populations of *C. diomedea* would still be in decline even in the absence of mortality due to fisheries bycatch or depredation (Genovart et al., 2017). The distribution of fishing vessels has been found to alter the foraging behaviour and movement patterns of shearwaters (e.g. *C. diomedea*), leading to greater probability of being caught as bycatch (Bartumeus et al., 2010), which is likely the main threat contributing to those intractable declines (Figure 2). Thus, the resilience of wide-ranging species, even among the same species, may be more nuanced and related to behavioural differences e.g., (Krüger et al., 2019), threat distributions, and localised foraging conditions (Bartumeus et al., 2010).

Larger seabirds have lower annual mortality thresholds, although this relationship is only weak. Indeed, body size explains only 9% of the annual mortality threshold (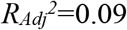; *t*=-2.8; *P*=0.02; Pagel’s λ=0.52, CI: 0.06-0.87, *P*<0.001; Fig. 3C; Table S3). Body size is generally a good indicator of extinction vulnerability for birds (Gaston & Blackburn, 1995), and for interactions with multiple marine threats. Larger birds have been found with greater loads of plastics (Ryan, 2016; Wilcox, Van Sebille, & Hardesty, 2015) and the demographic impact of fisheries bycatch is greater for albatrosses, large petrels, and shearwaters, due to low fecundity and high propensity to interact with fishing vessels (Richard et al., 2017; 2010). Other studies have shown large-bodied seabirds still declined following predator eradication, likely due to at-sea stressors (2010). All the species with high risk to marine threats in our analysis (*C. diomedea, C. borealis*, and *A. carneipes*) are large-bodied (all ~700 g). However, other larger bodied species such as *A. grisea* (Sooty shearwater) and *P. gouldi* have low risk ratios (Table 1; online material). Thus, inferring risk to marine threats from multiple shared traits may not be straightforward, particularly in the absence of depredation. Ecological and morphological traits may influence the tendency of a species to interact with a threat. Nonetheless, the spatial co-occurrence of threats and foraging opportunities, coupled with behaviour of individuals may exert more control over population level effects.

Our analyses suggest that gains from predator eradication are sufficient to offset mortality of a large proportion (82%) of seabird species from multiple marine threats of plastic ingestion, fisheries interactions, and climate change (Fig. 2; Table 1). However, for some species, predator eradication may not be a panacea (Figs. 1 & 2; Table 1). If at-sea adult mortality increases due to intensifying marine threats, then vulnerable species will continue to experience disproportionate declines (Genovart et al., 2017). For example, stable isotope analyses suggest either spatial shifts of, or reduced availability of prey indicating the potential for intensifying impacts from climate change (Bond & Lavers, 2014). This portends significant conservation challenges following predator eradication for seabirds highly impacted by marine threats (Croxall et al., 2012; Oro, 2014; Rodríguez et al., 2019; Wilcox et al., 2015).

While our phylogenetic analysis results align with the general consensus on vulnerability, and for risk in the colonies we analyse here, there are contradictions and species-specific responses to often ambiguous impacts of multiple marine threats. The uncertainty of inferring risk to multiple marine threats from closely-related species means that systematic population surveys after predator eradications are essential to detect colony recovery rates, and how threats may be affecting a population. The value of such monitoring is twofold; first, it can expediently inform managers of when additional conservation actions may be needed for species that fail to recover following predator eradication; and second, it can improve our ability to identify risk factors to marine threats for understudied species.

## Acknowledgements

Thanks to D.R. Towns, S. Avery-Gomm. SBB was supported by the AUT Pro-vice Chancellor’s Doctoral Scholarship and the David H. Smith Postdoctoral Research Fellowship; RS-G was supported by the Australian Research Council (DE140100505) and UK’s Natural Environment Research Council (R/142195-11-1). We thank J-D Lebreton and S. Devillard for providing us with data from BIDDABA. The authors declare no conflicts of interest.

## Author Contributions

SBB and HPJ developed the research question. SBB collected the data SBB, RSG, and YR did the analysis, SBB led the writing, and all authors contributed to writing and editing of the manuscript.

## References

Bærum, K. M., Anker-Nilssen, T., Christensen-Dalsgaard, S., Fangel, K., Williams, T., & Vølstad, J. H. (2019). Spatial and temporal variations in seabird bycatch: Incidental bycatch in the Norwegian coastal gillnet-fishery. PloS One, 14(3), e0212786.

Barbraud, C., Rolland, V., Jenouvrier, S., Nevoux, M., Delord, K., & Weimerskirch, H. (2012). Effects of climate change and fisheries bycatch on Southern Ocean seabirds: A review. Marine Ecology Progress Series, 454, 285–307. https://doi.org/10.3354/meps09616

Bartumeus, F., Giuggioli, L., Louzao, M., Bretagnolle, V., Oro, D., & Levin, S. A. (2010). Fishery discards impact on seabird movement patterns at regional scales. Current Biology, 20(3), 215–222.

Belda, E. J., & Sanchez, A. (2001). Seabird mortality on longline fisheries in the western Mediterranean: Factors affecting bycatch and proposed mitigating measures. Biological Conservation, 98(3), 357–363.

Birdlife International. (2016). Species Distribution data. Retrieved from http://datazone.birdlife.org/species/requestdis

Bond, A. L., & Lavers, J. L. (2014). Climate change alters the trophic niche of a declining apex marine predator. Global Change Biology, 20(7), 2100–2107.

Borrelle, S. B., Boersch-Supan, P. H., Gaskin, C. P., & Towns, D. R. (2016). Influences on recovery of seabirds on islands where invasive predators have been eradicated, with a focus on Procellariiformes. Oryx, 52(2), 346–358.

Bost, C. A., Cotté, C., Terray, P., Barbraud, C., Bon, C., Delord, K., … Guinet, C. (2015). Large-scale climatic anomalies affect marine predator foraging behaviour and demography. Nature Communications, 6, 8220.

Brooke, M. de L., Bonnaud, E., Dilley, B. J., Flint, E. N., Holmes, N. D., Jones, H. P., … Surman, C. (2018). Seabird population changes following mammal eradications on islands. Animal Conservation, 21(1), 3–12.

Burthe, S. J., Wanless, S., Newell, M. A., Butler, A., & Daunt, F. (2014). Assessing the vulnerability of the marine bird community in the western North Sea to climate change and other anthropogenic impacts. Marine Ecology Progress Series, 507, 277–295.

Buxton, R. T., Jones, C. J., Moller, H., & Towns, D. R. (2014). Drivers of Seabird Population Recovery on New Zealand Islands after Predator Eradication. Conservation Biology, 28(2), 333–344.

Clukey, K. E., Lepczyk, C. A., Balazs, G. H., Work, T. M., Li, Q. X., Bachman, M. J., & Lynch, J. M. (2018). Persistent organic pollutants in fat of three species of Pacific pelagic longline caught sea turtles: Accumulation in relation to ingested plastic marine debris. Science of the Total Environment, 610, 402–411.

Cooke, R. S., Eigenbrod, F., & Bates, A. E. (2019). Projected losses of global mammal and bird ecological strategies. Nature Communications, 10(1), 2279.

Croxall, J. P., Butchart, S. H. M., Lascelles, B., Stattersfield, A. J., Sullivan, B., Symes, A., & Taylor, P. (2012). Seabird conservation status, threats and priority actions: A global assessment. Bird Conservation International, 22(01), 1–34.

Day, R. H., Wehle, D. H., & Coleman, F. C. (1985). Ingestion of plastic pollutants by marine birds. 2, 34. US Dep. Commer. NOAA Tech Memo NMFS Honolulu, Hawaii.

Dillingham, P. W., Moore, J. E., Fletcher, D., Cortés, E., Alexandra Curtis, K., James, K. C., & Lewison, R. L. (2016). Improved estimation of intrinsic growth rmax for long-lived species: Integrating matrix models and allometry. Ecological Applications, 26(1), 322–333.

Freckleton, R. P., Harvey, P. H., & Pagel, M. D. (2002). Phylogenetic analysis and comparative data: A test and review of evidence. American Naturalist, 160(6), 712–726. Retrieved from http://www.jstor.org/stable/10.1086/343873

Frederiksen, M., Harris, M. P., Daunt, F., Rothery, P., & Wanless, S. (2004). Scale-dependent climate signals drive breeding phenology of three seabird species. Global Change Biology, 10(7), 1214–1221.

Gaston, K. J., & Blackburn, T. M. (1995). Birds, body size and the threat of extinction. Philosophical Transactions of the Royal Society B: Biological Sciences, 347(1320), 205–212.

Genovart, M., Doak, D. F., Igual, J., Sponza, S., Kralj, J., & Oro, D. (2017). Varying demographic impacts of different fisheries on three Mediterranean seabird species. Global Change Biology.

Giudici, A., Navarro, J., Juste, C., & González-Solís, J. (2010). Physiological ecology of breeders and sabbaticals in a pelagic seabird. Journal of Experimental Marine Biology and Ecology, 389(1), 13–17.

Greene, B. S., Taylor, G. A., & Earl, R. (2015). Distribution, population status and trends of grey-faced petrel (Pterodroma macroptera gouldi) in the northern North Island, New Zealand. Notornis, 62(3), 143–161.

Ismar, S. M., Baird, K. A., Gaskin, C. P., Taylor, G. A., Tennyson, A. J., Rayner, M. J., … Imber, M. J. (2014). A case of natural recovery after the removal of invasive predators–community assemblage changes in the avifauna of Burgess Island. Notornis, 61, 188–195.

IUCN. (2017). IUCN Red List of Threatened Species^TM^ (Vol. 2017). Retrieved from http://www.iucnredlist.org

Jetz, W., Thomas, G., Joy, J., Hartmann, K., & Mooers, A. (2012). The global diversity of birds in space and time. Nature, 491(7424), 444.

Jones, H. P., Holmes, N. D., Butchart, S. H. M., Tershy, B. R., Kappes, P. J., Corkery, I., … Burbidge, A. A. (2016). Invasive mammal eradication on islands results in substantial conservation gains. Proceedings of the National Academy of Sciences, 201521179.

Jones, H. P., Tershy, B. R., Zavaleta, E. S., Croll, D. A., Keitt, B. S., Finkelstein, M. E., & Howald, G. R. (2008). Severity of the effects of invasive rats on seabirds: A global review. Conservation Biology, 22(1), 16–26.

Joseph, L., & Bélisle, P. (2012). Gamma parms from quantiles. Retrieved from http://www.medicine.mcgill.ca/epidemiology/joseph/pbelisle/GammaParmsFromQuantiles.html

Kappes, P. J., & Jones, H. P. (2014). Integrating seabird restoration and mammal eradication programs on islands to maximize conservation gains. Biodiversity and Conservation, 23(2), 503–509.

Krüger, L., Pereira, J. M., Paiva, V. H., & Ramos, J. A. (2019). Personality influences foraging of a seabird under contrasting environmental conditions. Journal of Experimental Marine Biology and Ecology, 516, 123–131.

Lavers, J. L., Bond, A. L., & Hutton, I. (2014). Plastic ingestion by flesh-footed shearwaters (Puffinus carneipes): Implications for fledgling body condition and the accumulation of plastic-derived chemicals. Environmental Pollution, 187, 124–129.

Lavers, J. L., Wilcox, C., & Donlan, C. J. (2010). Bird demographic responses to predator removal programs. Biological Invasions, 12(11), 3839–3859.

Lawler, J. J., Campbell, S. P., Guerry, A. D., Kolozsvary, M. B., O’Connor, R. J., & Seward, L. C. (2002). The scope and treatment of threats in endangered species recovery plans. Ecological Applications, 12(3), 663–667.

Niel, C., & Lebreton, J.-D. (2005). Using demographic invariants to detect overharvested bird populations from incomplete data. Conservation Biology, 19(3), 826–835.

Orme, D. (2013). The caper package: Comparative analysis of phylogenetics and evolution in R. R Package Version, 5(2).

Oro, D. (2014). Seabirds and climate: Knowledge, pitfalls, and opportunities. Frontiers in Ecology and Evolution, 2, 79.

Paniw, M., Ozgul, A., & Salguero-Gómez, R. (2018). Interactive life-history traits predict sensitivity of plants and animals to temporal autocorrelation. Ecology Letters, 21(2), 275–286.

Paradis, E., Claude, J., & Strimmer, K. (2004). APE: analyses of phylogenetics and evolution in R language. Bioinformatics, 20(2), 289–290.

Provencher, J., Borrelle, S. B., Sherley, R. B., Avery-Gomm, S., Hodum, P. J., Bond, A. L., … Mallory, M. L. (2018). Seabirds. In World Seas, Volume III: Ecological Issues and Environmental Impacts. Cambridge, MA, USA: Elsevier, Inc.

Provencher, J., Gaston, A. J., Mallory, M. L., O’hara, P. D., & Gilchrist, H. G. (2010). Ingested plastic in a diving seabird, the thick-billed murre (Uria lomvia), in the eastern Canadian Arctic. Marine Pollution Bulletin, 60(9), 1406–1411.

R Core Team. (2013). R: A language and environment for statistical computing (R. F. for S. Computing, Ed.). Vienna, Austria.

Revell, L. J. (2012). phytools: An R package for phylogenetic comparative biology (and other things). Methods in Ecology and Evolution, 3(2), 217–223.

Richard, Y., & Abraham, E. R. (2013). Application of Potential Biological Removal methods to seabird populations (Vol. 6480).

Richard, Y., Abraham, E. R., & Filippi, D. (2017). Assessment of the risk of commercial fisheries to New Zealand seabirds, 2006-07 to 2014-15. Ministry for Primary Industries, Manatū Ahu Matua.

Riou, S., Gray, C. M., Brooke, M. de L., Quillfeldt, P., Masello, J. F., Perrins, C., & Hamer, K. C. (2011). Recent impacts of anthropogenic climate change on a higher marine predator in western Britain. Marine Ecology Progress Series, 422, 105–112.

Rodríguez, A., Arcos, J. M., Bretagnolle, V., Dias, M. P., Holmes, N. D., Louzao, M., … Rodríguez, B. (2019). Future Directions in Conservation Research on Petrels and Shearwaters. Frontiers in Marine Science, 6, 94.

Roman, L., Bell, E., Wilcox, C., Hardesty, B. D., & Hindell, M. (2019). Ecological drivers of marine debris ingestion in Procellariiform Seabirds. Scientific Reports, 9(1), 916.

Rowe, S. (2010). Level 1 Risk Assessment for incidental seabird mortality associated with New Zealand fisheries in the NZ-EEZ. Marine Conservation Services, Department of Conservation, Wellington.

Ryan, P. G. (1987). The incidence and characteristics of plastic particles ingested by seabirds. Marine Environmental Research, 23(3), 175–206.

Ryan, P. G. (2016). Ingestion of plastics by marine organisms.

Sæther, B.-E., & Engen, S. (2010). Population analyses. In A. P. Møller, W. Fiedler, & P. Berthold (Eds.), Effects of climate change on birds. Oxford, UK: Oxford University Press.

Salguero-Gómez, R., Jones, O. R., Archer, C. R., Bein, C., de Buhr, H., Farack, C., … Hoppe, G. (2016). COMADRE: a global data base of animal demography. Journal of Animal Ecology, 85(2), 371–384.

Schreiber, E. A., & Burger, J. (2002). Biology of Marine Birds. CRC Press Boca Raton, Florida.

Sutherland, W. J., Bellingan, L., Bellingham, J. R., Blackstock, J. J., Bloomfield, R. M., Bravo, M., … Cohen, A. S. (2012). A collaboratively-derived science-policy research agenda. PloS One, 7(3), e31824.

Symonds, M. R. E., & Blomberg, S. P. (2014). A Primer on Phylogenetic Generalised Least Squares. In L. Z. Garamszegi (Ed.),. Modern Phylogenetic Comparative Methods and Their Application in Evolutionary Biology (pp. 105–130).

Tanaka, K., Takada, H., Yamashita, R., Mizukawa, K., Fukuwaka, M., & Watanuki, Y. (2015). Facilitated leaching of additive-derived PBDEs from plastic by seabirds’ stomach oil and accumulation in tissues. Environmental Science & Technology.

Towns, D. R., Byrd, G. V., Jones, H. P., Rauzon, M. J., Russell, J. C., & Wilcox, C. (2011). Impacts of introduced predators on seabirds. Seabird Islands. Ecology, Invasion, and Restoration. Oxford University Press, New York, 56–90.

van Franeker, J. A., & Law, K. L. (2015). Seabirds, gyres and global trends in plastic pollution. Environmental Pollution, 203, 89–96.

Votier, S. C., Hatchwell, B. J., Mears, M., & Birkhead, T. R. (2009). Changes in the timing of egg-laying of a colonial seabird in relation to population size and environmental conditions. Marine Ecology Progress Series, 393, 225–233.

Warham, J. (1990). The petrels: Their ecology and breeding systems. A&C Black.

Weimerskirch, H. (2001). Seabird demography and its relationship with the marine environment. In E. A. Schreiber & J. Burger (Eds.), Biology of Marine Birds (EA Schreiber and J. Burger, Eds.), CRC Press, Boca Raton, FL (pp. 115–136).

Wilcox, C., Van Sebille, E., & Hardesty, B. D. (2015). Threat of plastic pollution to seabirds is global, pervasive, and increasing. Proceedings of the National Academy of Sciences of the United States of America, 112(38), 11899–11904.

