## Supplementary materials for "Not all seabird species can overcome marine threats when predator removal at their colonies is prioritised"

**This PDF file includes:**

Supplementary text

Figs. S1 to S7

Tables S1 to S4

References for SI reference citations

Datasets S1 – S2

### Methods

Parameter estimates for the demographic analysis involving 81 species were from the primary literature, and renowned grey literature (*e.g.*, Birdlife International, and NZ Birds Online websites). Values for α and *s* were taken as given in BIDDABA (n=13 species); for the six species from COMADRE, α was calculated with methods described in Caswell (Caswell, 2001 p. 124), whereas *s* was calculated as the mean column-sums of the sub-matrix of survival-dependent transitions corresponding to adult reproductive stages, weighted by its stable stage distribution (see Salguero-Gómez et al., 2016). The IUCN Red List threatened species ranking and global population size data were obtained from the IUCN Red List (IUCN, 2017).

The dataset of colonies on islands where predators were removed were selected because there were estimates of the population size, which are needed for the Demographic Invariant Method (DIM). The regional population estimates were obtained from The Northern New Zealand Seabird Trust (Unpublished; Table S2) and from Brooke et al. (Brooke et al., 2018). We excluded colonies that were smaller than 50 individuals because smaller colonies are thought to be more prone to extinction, because they are more affected my demographic stochasticity, which may be related to minimum viable population sizes (Caughley, 1994).

The ecological and morphometric information of each species was used to correlate key attributes of the species niche to its vulnerability to at-sea threats. These include foraging strategy (*i.e.*, the primary method of feeding, such as pursuit diving, surface seizing, pattering), primary prey type (i.e., cephalopods, fish) collated from Ashmole (1971), Schreiber and Burger (Schreiber & Burger, 2002), Del Hoyo et al. (2011), and NZ BirdsOnline. Mean (female) adult body mass (B) data were obtained from the CRC

Handbook of Avian Mass files (Dunning, 2013). No uncertainty for body mass estimates were included in the model. At-sea distribution data were from Birdlife International (2016). We used taxonomy and nomenclature from Birdlife International (Birdlife International, 2014), which differed for some species in Jetz et al. (2012) that was used for the phylogenetic analysis. All parameters used in the risk analysis model for 37 colonies of 17 species can be found in Dataset S1, and parameters for the trait analysis including 81 procellariiform species in Dataset S2.

**Data quality**

Seabirds breed on remote islands, therefore collecting accurate demographic data consistently can be logistically and financially challenging, resulting in large uncertainty of parameter estimates (Richard & Abraham, 2013a). Sources of bias and error in the demographic parameters may stem from multiple factors. For example, the estimates of adult survival for most species are likely to underestimate natural rates. This is because it is impossible to remove the effect of anthropogenic sources of mortality from studies that these values are derived from. A brief summary of our approach for data collation and discussion of the limitations is detailed as follows:

**Adult survival (*s*):** While most seabirds have an adult survival of > 90% in optimal conditions, those with values at or below 90%, optimal adult survival rates would be expected to be ~95% (Richard & Abraham, 2013a). Therefore, population growth rates using the demographic invariant method (DIM) are likely to be underestimated. For the risk analysis of 17 species we used the highest value of adult survival reported in the literature, with the assumption that this estimate was likely to be closest to adult survival in the absence of predators, given the available habitat and absence of introduced predators on the islands (Borrelle, Boersch-Supan, Gaskin, & Towns, 2016). This approach allowed us to estimate the potential impacts to a population specifically from the three marine threats evaluated. Where data on adult survival for a species were unknown (n=3) we assumed that adult survival was 0.93 SE ± 0.03 (Brooke et al., 2010).

**Age of first reproduction (a)** We used the mean value of age of first reproduction (α). Age of first reproduction data may be estimated from a small sample size, leading to either an over- or under-estimation. In such cases, the population growth rate ($\lambda_{max}$) will be over or underestimated. Uncertainties in α were incorporated into model outputs through a bootstrapping process with 1000 iterations from the gamma distribution, and expressed as the mean and standard deviation of the mean.

**Population size and demographic parameters:** Estimates of the population size of most species is embedded with bias and uncertainty. This is because many of the population surveys come from data older than eight years, and in most cases, there is a paucity of details about survey methods. Population estimates may come from a one off survey, which may have been a good or bad year for individuals choosing to breed (Frederiksen, Harris, Daunt, Rothery, & Wanless, 2004), thereby over- or under-estimating breeding pair numbers. In addition, these counts may not accurately count non-breeding birds (*i.e.*, immature individuals, or those on sabbatical). For the phylogenetic comparative analysis. we used the minimum *total* population estimates as per the IUCN Red List and Birdlife International (Birdlife International, 2016; IUCN, 2017). When only estimates of breeding pairs were available, we multiplied the breeding pair estimate by 2.3, which includes ~30% for non-breeders (*n=2)* (Cornell Lab of Ornithology, 2019). Because of the uncertainty in estimates of the population size, we used a bootstrapping process with 5000 iterations from a log-normal distribution, and expressed as the mean output and standard deviation of the mean. To account for the uncertainty in population estimates, we assigned all population estimate a standard deviation of 0.05.

We did not account for density dependence in both positive and negative directions when calculating $\lambda_{max}$. Negative density dependence is where $\lambda$ declines as the population density increases, in this instance it is assumed that fecundity is higher at low densities and decays at a constant rate (Morris & Doak, 2002). Although rare, the opposite can also occur, where rapid declines in fecundity occur at low population densities, and then increase at higher densities, given no limiting factors (*e.g.,* no resource or habitat limitations). The magnitude of negative density dependence across a range of densities can vary due to external or internal population level effects (Morris & Doak, 2002, p 38). Positive density dependence, sometimes referred to as Allee effects, lead to an increase in population growth rate as the population increases. Such responses are likely when resources are not limited, mating success is improved, group defense reduces predation (Morris & Doak, 2002).

Determining density dependence extent or type in a population in either direction is inherently difficult due to data limitations (Morris & Doak, 2002). Because the current knowledge of processes and effects on population growth rates is lacking, given the long-term and detailed studies required, there is uncertainty in estimating how density dependence may be affecting our results. Finally, we did not address the potential carrying capacity of the population, and assumed that the length of time to reach this point for most populations is beyond current temporal management plans (e.g. 200 years; 18), also that other threats or changes in levels of mortality from threats (e.g. climate change) will adjust the results of the model.

**Marine threat Data**

Detecting at-sea mortality is challenging because seabirds are scattered widely across their foraging ranges, and carcasses may float just below the surface, sink or be consumed by predators (Laist, 1997). Furthermore, in the incidence of birds being entangled in fishing ling or ropes may be mistaken for fisheries related bycatch – where animals are incidentally caught in active fishing gear, or in some cases in ghost nets rather than from mortality from plastic ingestion, although this is likely a very small proportion (Laist, 1997). On land many species are understudied, and land-based surveys provide no indication of the number of at-sea mortalities. In this paper, we address only the impacts of plastic pollution, climate change and commercial fisheries to our seabird populations. We acknowledge that our model does not include the full suite of marine threats that seabirds are exposed including such as disease, oil-spills, water-bound contaminants, hunting ((for comprehensive reviews of the full suite of threats to seabirds see; Provencher et al., 2018; and Rodríguez et al., 2019)). The methods used to estimate the potential fatalities from each of the marine threats included in our analysis are as follows:

**Fisheries:** We used the mean annual potential mortality from fisheries from Richard et al (Richard, Abraham, & Filippi, 2017) for 12 of the seabird species included in the colony analysis (Appendix 2). Richard et al.'s (22) estimate assumes that all birds killed in the fisheries were adults (98% of the necropsied birds were adults). The estimates for fisheries related mortality reported here do not account for mortality associated with international (beyond the EEZ), illegal and unregulated, or recreational fisheries, which may present a significant source of mortality for some species. Estimates for *Calonectris diomedea* were from Belda & Sanchez (Belda & Sanchez, 2001) and we used the same estimate for the closely related species *C. borealis*, which may underestimate the impact of fisheries bycatch for this species (Dataset 1). The species *Pterodroma hypoleuca, P. ultima* and *Puffinus puffinus* were assumed to be low risk from fisheries because are not highly reported as bycatch in the literature (IUCN, 2017). These three species were assumed to have 0.1% of the population killed by fisheries. The estimation of at-sea mortality due to a particular threat typically results in a high degree of imprecision. For example, when estimates of adult mortality in a fishery are reported, they are often calculated from a small number (typically < 10%) of ship-board observations (Richard et al., 2011). In addition, there are a lack of data on cryptic mortalities in commercial fishing operations, that is, birds that are killed may not be bought back on board the ship, may fail to be reported when the observer is off-duty, or not seen by the observer. Specific information on the relation between observed captures and total fatalities needs to be improved in order to improve the reliability of our risk assessment (Richard & Abraham, 2013c). Therefore, it is possible that our model will either fail to adequately quantify the risk of at-sea threats to a seabird species, or will classify species as being at risk when in they may not be.

**Plastic ingestion:** The physiological effects of plastic debris ingestion on seabirds may include; internal and external wounds, skin lesions and ulcerating sores, ingestion causing general debilitation, inhibiting feeding capacity, eventually leading to starvation, reductions in reproductive capacity, drowning, and impairment of predator avoidance (Auman, Ludwig, Giesy, & Colborn, 1997; Ryan, 1987; Vannela, 2012). We used the proportion of adults reported in the literature to ingest plastic – the frequency of occurrence - and estimated the proportion of the population on the islands affected (Avery-Gomm et al. *in review)*. We assumed that the colonies would be affected at the same rate as the frequency of occurrence reported in the literature (Appendix 2), and that of the proportion that ingest plastic, 0.5% of the affected population would die as a direct result of plastic ingestion. This approach may over- or under-estimate the impact of plastic ingestion on adult mortality as there is a lack of understanding about plastic retention in animals and what the long-term impacts may be on adult survival and populations (Rochman et al., 2016; Ryan, 2016). We tested the sensitivity of each of the species and colonies to plastic ingestion related mortality at 1% and 5% of the proportion of a population that is expected to ingest plastic (Figure S3 & S4).

**Climate Change:** Despite impressive research efforts that indicate seabirds are the most vulnerable group of avian fauna to climatic changes (Jenouvrier, 2013; Oro, 2014), there is high uncertainty in our model to predict adult mortality from climate change pressures. This is due to the difficulty in quantifying adult mortality directly due to the complex interactions affecting prey distributions and abundance (Oro, 2014; Sæther & Engen, 2010), confounded by the lack of published studies on the effects of climate change for the 17 species included in our marine threats risk analysis. The influence of climatic changes on seabird populations may exert either positive or negative changes to a population in response to resource availability and distribution, breeding phenology or impacts on habitat (Engen & Sæther, 2016; Jenouvrier, 2013). In addition, other factors, such as density dependence, inter- and intra-specific competition, and scale dependent variability in climatic stressors will influence how an individual or population will respond (Jenouvrier, 2013; Oro, 2014). Thus, attributing changes to adult survival directly to a specific climate driver is complex, and generalising among species can lead to erroneous assumptions (Oro, 2014). Until reliable estimates adult mortality from anthropogenic marine threats to seabirds exist, accurately estimating the population-level effects on seabirds will remain challenging. Because of the uncertainty in estimating the level of mortality caused by climate change, we estimated the impact of climate change as causing 0.5% mortality in a population and tested the sensitivity to risk of 1% and 5% adult mortality for each of the 37 colonies in our risk analysis (Figs. S5 & S6).

### Extended results

**Demographic results for 37 colonies of 17 species on islands eradicated of invasive predators.**

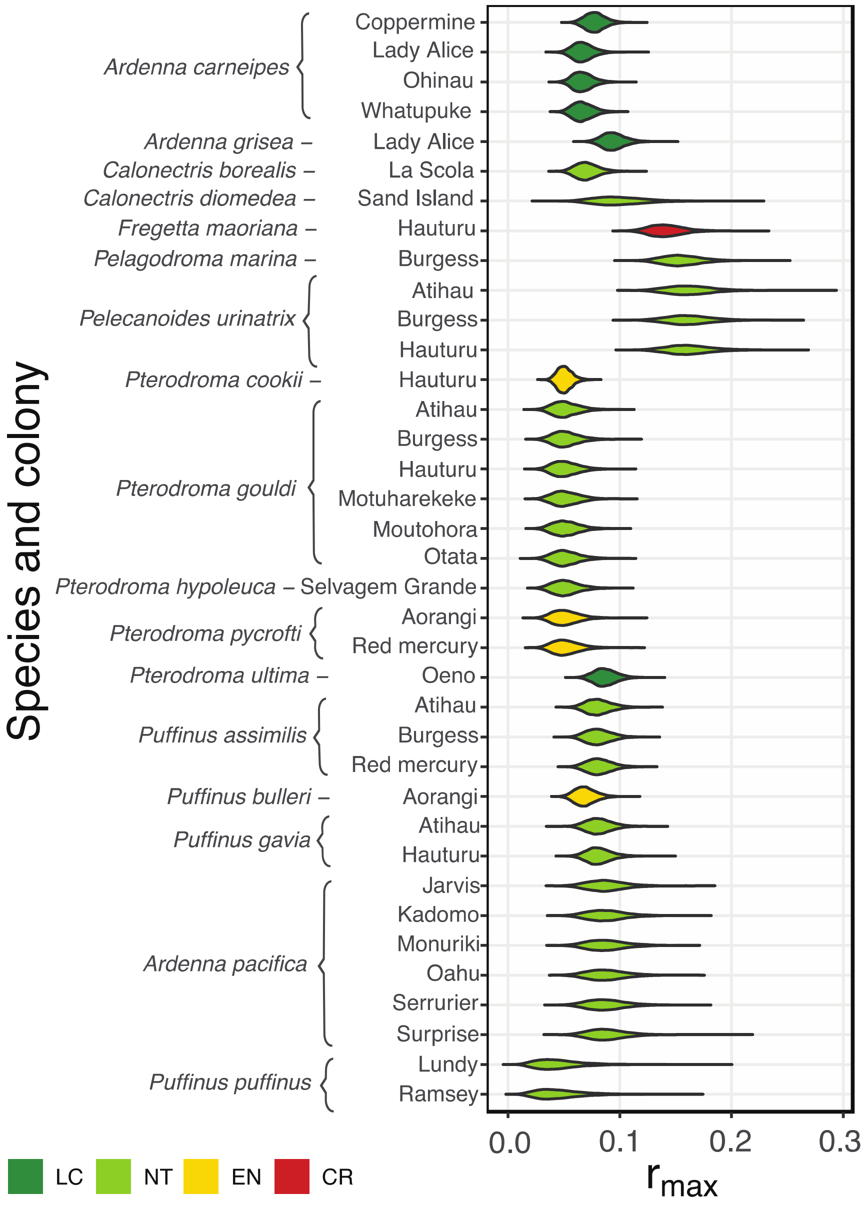


**Figure S1:** The model calculated optimal population growth rate ($r_{max}$) for each of the 37 colonies, on 24 islands where invasive predators have been eradicated, including 17 species. Colors for each species correspond to the IUCN Red List status: LC Least Concern in dark green; NT Near threatened in light green; VU Vulnerable in yellow; CR Critically Endangered in red.

**Impact of marine threats on the recovery of seabird species post-eradication**

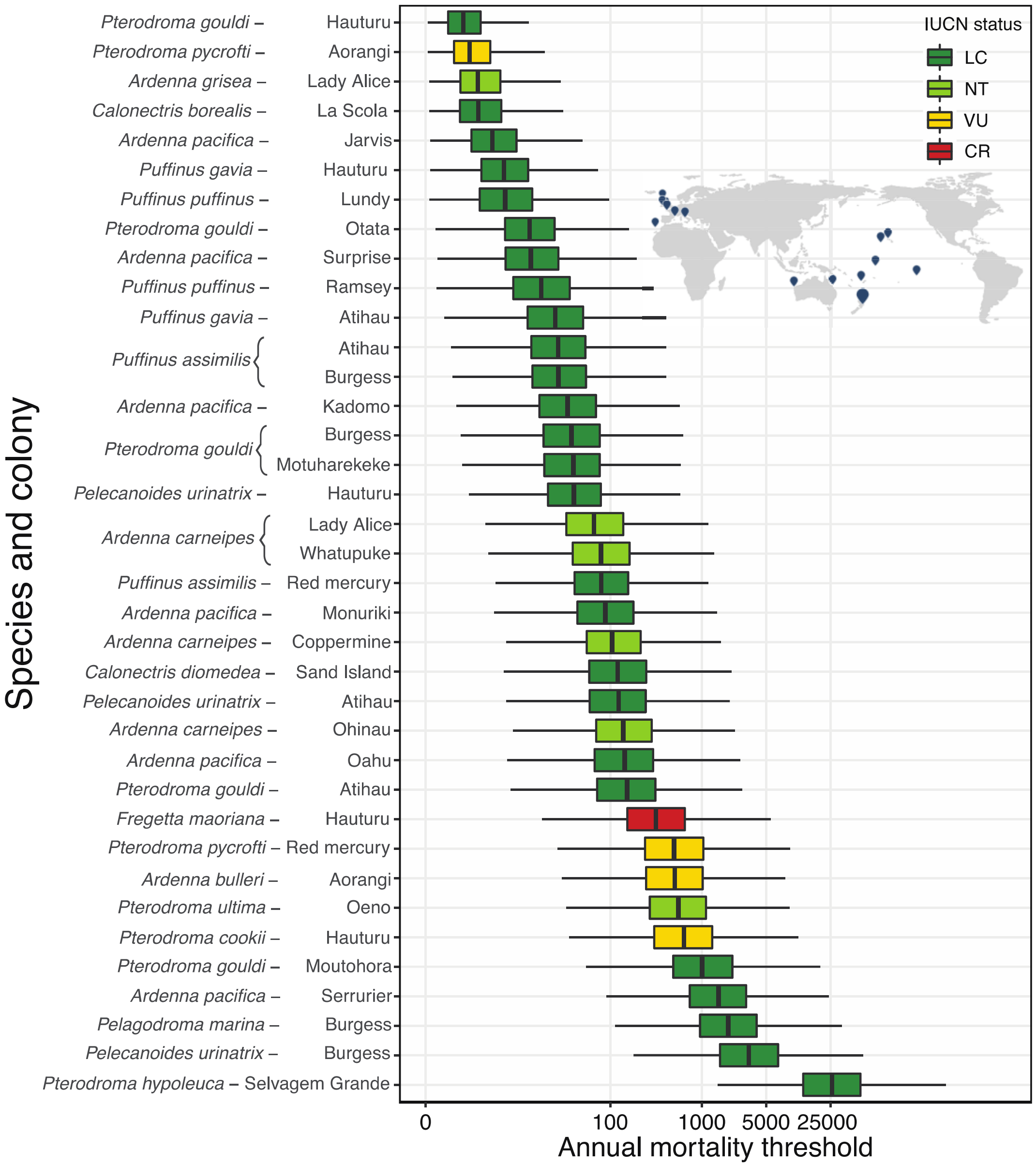


**Figure S2:** The model calculated annual mortality threshold for 17 species from 37 colonies on 24 islands where invasive predators have been eradicated (map inset**)**. The annual mortality threshold for each colony is ranked from lowest (top) to highest. Colors for each species correspond to the IUCN Red List status: LC Least Concern in dark green; NT Near threatened in light green; VU Vulnerable in yellow; CR Critically Endangered in red.

**Plastic ingestion related mortality risk ratio sensitivity analysis:**

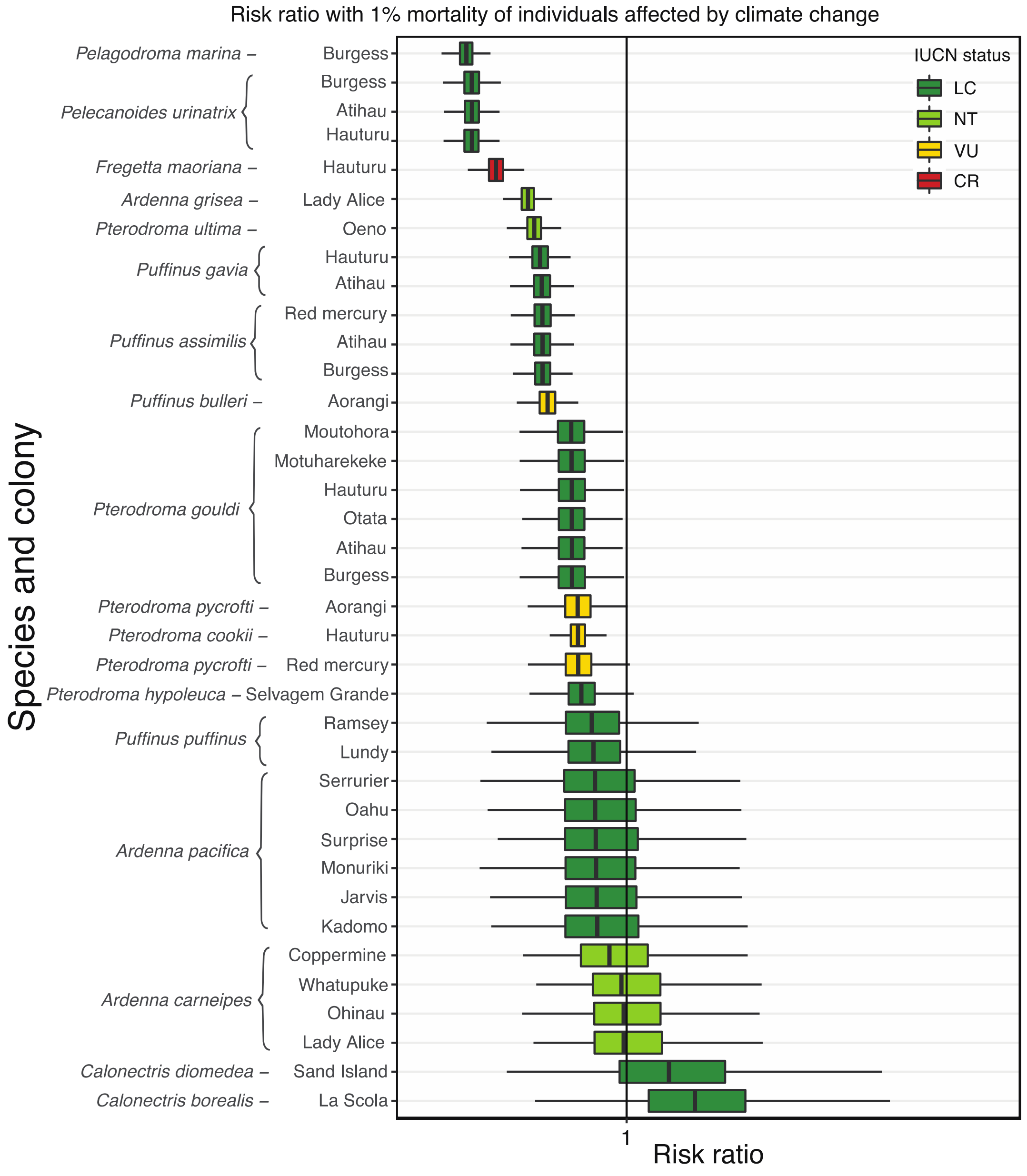


**Figure S3:** Sensitivity analysis showing the risk for each of the 37 colonies for the 17 species to plastic ingestion mortality at 1% for the proportion of individuals affected (Appendix 1). The risk ratio was calculated as potential mortalities yr-1 / annual mortality threshold (Richard & Abraham, 2013b); when this risk ratio ≥1, adult mortality from each of the evaluated threats may impede the recovery of a colony even after predator eradication. Colors for each species correspond to the IUCN Red List status: LC Least Concern in dark green; NT Near threatened in light green; VU Vulnerable in yellow; CR Critically Endangered in red.

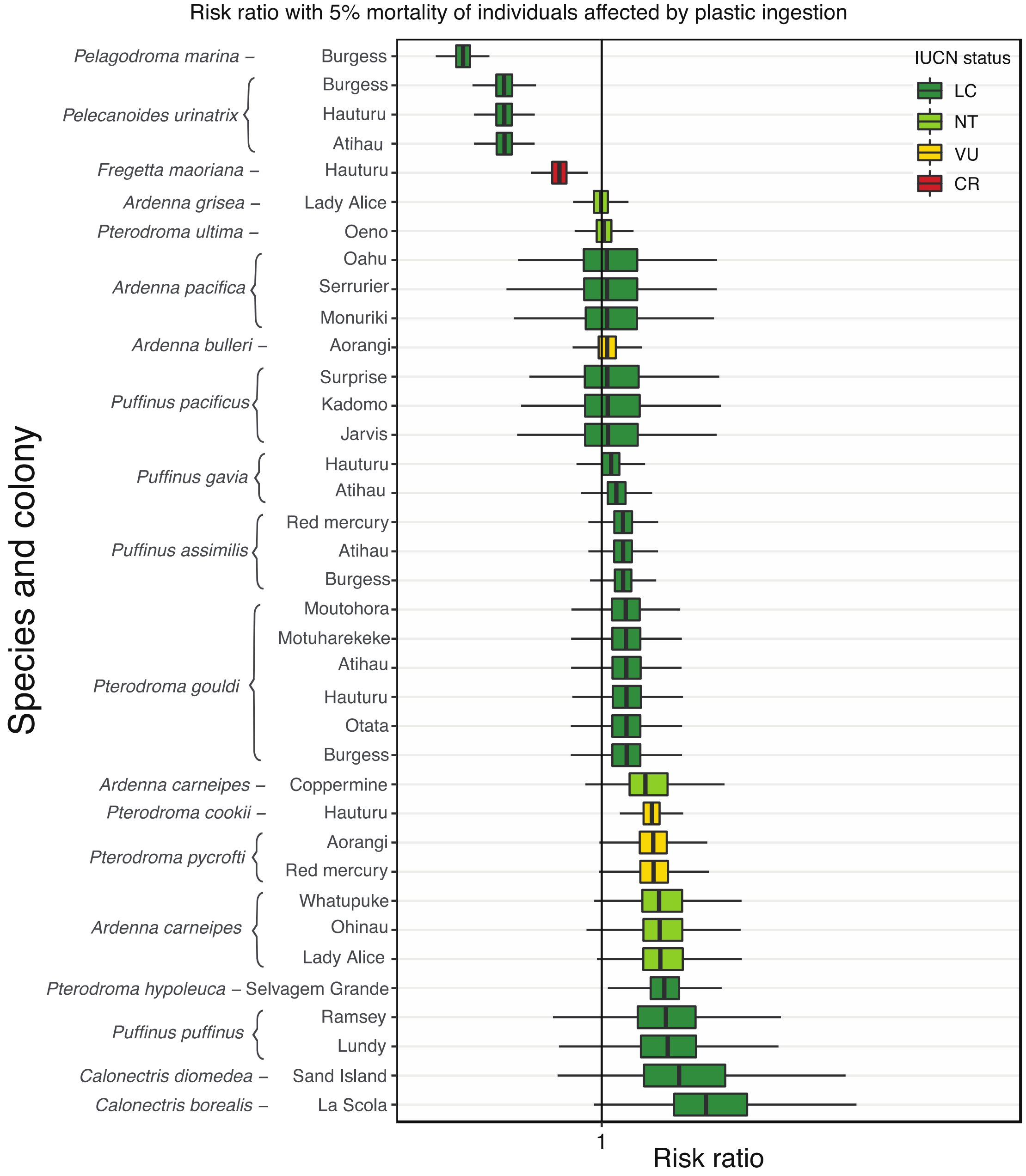


**Figure S4:** Sensitivity analysis showing the risk for each of the 37 colonies for the 17 species to plastic ingestion mortality at 5% for the proportion of individuals affected. The risk ratio was calculated as potential mortalities yr-1 / annual mortality threshold (Richard & Abraham, 2013b); when this risk ratio ≥1, adult mortality from each of the evaluated threats may impede the recovery of a colony even after predator eradication. Colors for each species correspond to the IUCN Red List status: LC Least Concern in dark green; NT Near threatened in light green; VU Vulnerable in yellow; CR Critically Endangered in red.

**Climate change risk ratio sensitivity analysis:**

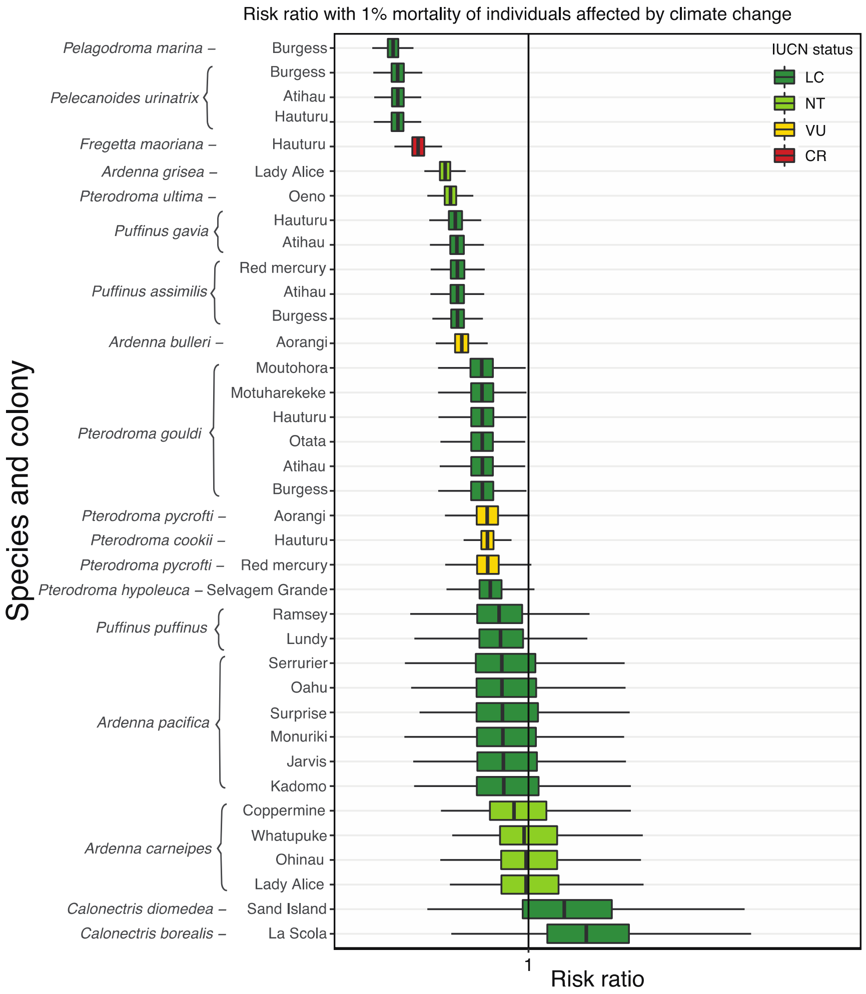


**Figure S5:** Sensitivity analysis showing the risk for each of the 37 colonies for the 17 species to climate change mortality at 1%. The risk ratio was calculated as potential mortalities yr-1 / annual mortality threshold (Richard & Abraham, 2013b); when this risk ratio ≥1, adult mortality from each of the evaluated threats may impede the recovery of a colony even after predator eradication. Colors for each species correspond to the IUCN Red List status: LC Least Concern in dark green; NT Near threatened in light green; VU Vulnerable in yellow; CR Critically Endangered in red.

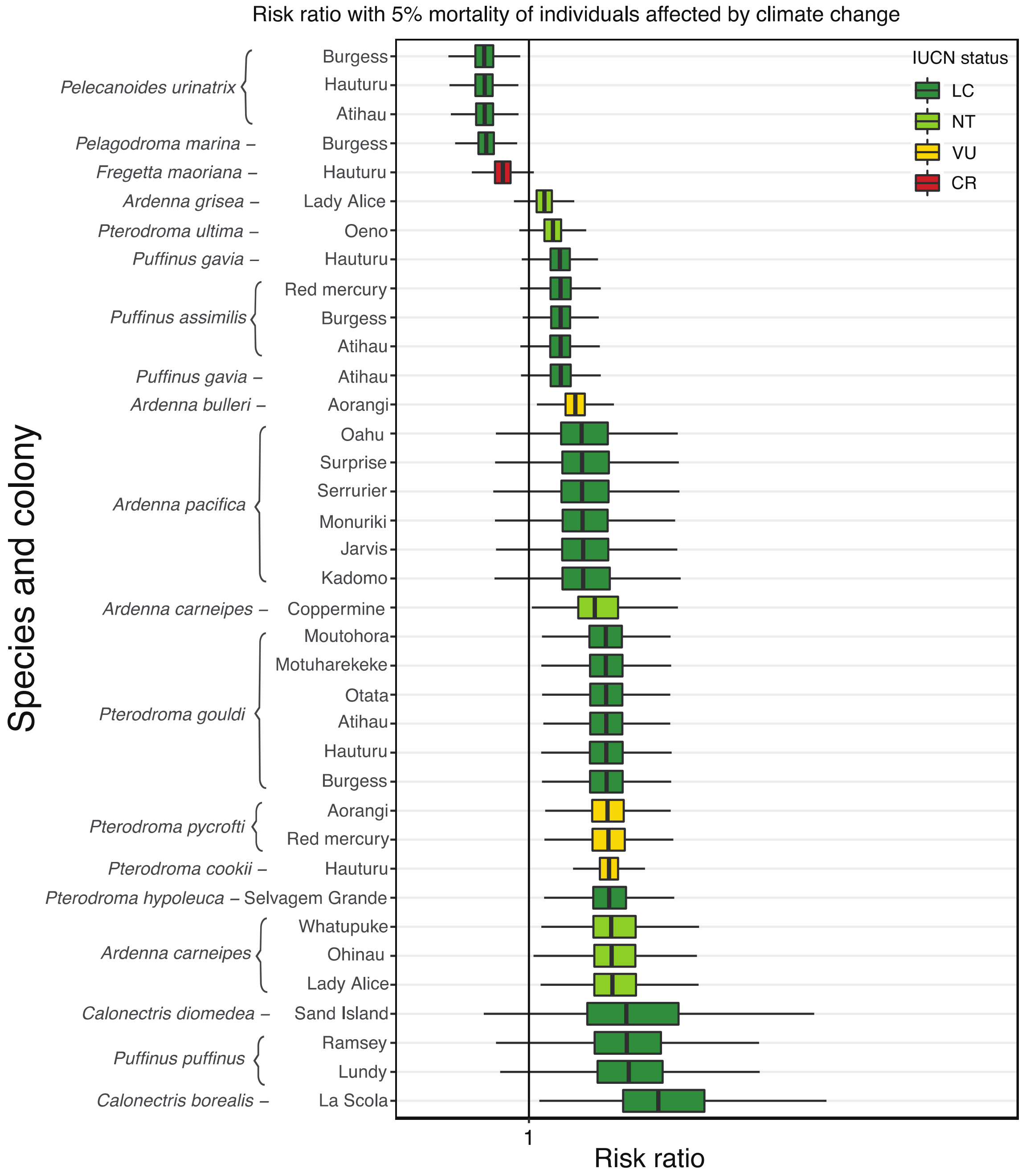


**Figure S6:** Sensitivity analysis showing the risk for each of the 37 colonies for the 17 species to climate change mortality at 5%. The risk ratio was calculated as potential mortalities yr-1 / annual mortality threshold (Richard & Abraham, 2013b); when this risk ratio ≥1, adult mortality from each of the evaluated threats may impede the recovery of a colony even after predator eradication. Colors for each species correspond to the IUCN Red List status: LC Least Concern in dark green; NT Near threatened in light green; VU Vulnerable in yellow; CR Critically Endangered in red.

**Annual mortality threshold and trait analysis for 81 procellariiform seabirds.**

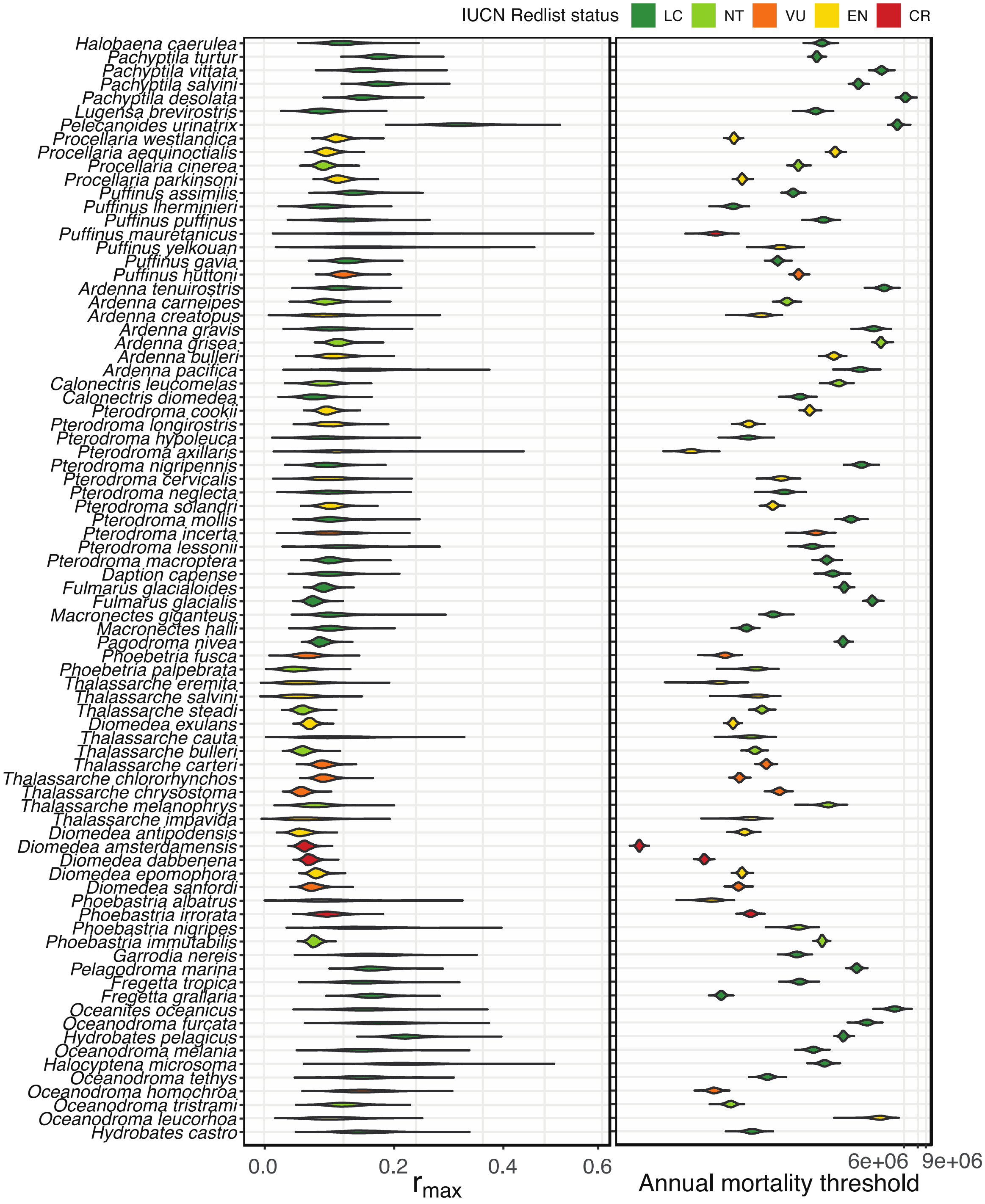


**Figure S7:** Annual population growth rate ($r_{max}$) and potential annual mortality threshold for the 81 seabird species included in the phylogenetic comparative analysis to investigate if shared traits can inform risk to marine threats.

**Table S2.** Annual mortality threshold model validation Tukey's Test results (p=0.0001). See Main text; Figure 2A.

| IUCN Category | Means | Group |
|---|---|---|
| Least Concern (LC) | 10.93 | a |
| Near Threatened (NT) | 9.32 | ab |
| Vulnerable (VU) | 7.93 | bc |
| Endangered (EN) | 7.68 | bc |
| Critical (CR) | 4.76 | c |

**Table S3.** Species traits ecological traits of at-sea distribution, adult body size, and foraging strategies of pursuit diving and surface filtering predict a species' annual mortality threshold (in bold). The annual mortality threshold is the limit of individuals in a population that can be 'harvested' for the population to remain stable or increase. The phylogenetic generalised least squares models below present the relative influence of the foraging strategies of surface filtering, and pursuit diving, and morphometric variables (adult mean body size) on the annual mortality threshold for the 81 seabird species examined. Pagel's $\lambda$ is a proxy to phylogenetic signal, with values close to 1 indicating high phylogenetic inertia (i.e. trait under consideration is highly preserved in the species pool). Non- significant results are shown.

| Model | t-statistic (df=79) | Pr | $R_{Adj}^2$ | Pagel's λ |
|---|---|---|---|---|
| **AMT_mean ~ surface_filtering** | **5.02** | **2.20E-05** | **0.24** | **1.00E-06** |
| **AMT_mean ~ pursuit_diving** | **4.20** | **0.0004** | **0.18** | **0.42** |
| **AMT_log ~ range_log** | **3.66** | **0.002** | **0.14** | **0.742** |
| **AMT_log ~ biomass_log** | **-2.82** | **0.025** | **0.091** | **0.42** |
| AMT_mean ~ scavenging | -1.87 | 0.169 | 0.042 | 0.48 |
| AMT_mean ~ surface_seizing | -1.63 | 0.261 | 0.032 | 0.54 |
| AMT_mean ~ pattering | -1.27 | 0.398 | 0.020 | 0.54 |
| AMT_mean ~ fishfood | -0.99 | 0.573 | 0.012 | 0.55 |
| AMT_mean ~ crustaceans | 0.82 | 0.676 | 0.008 | 0.54 |
| AMT_mean ~ pursuit_plunging | 0.78 | 0.688 | 0.007 | 0.56 |
| AMT_mean ~ plunging | 0.62 | 0.788 | 0.005 | 0.56 |
| AMT_mean ~ dipping | 0.47 | 0.819 | 0.003 | 0.55 |
| AMT_mean ~ cephalopods | -0.41 | 0.819 | 0.002 | 0.56 |
| AMT_mean ~ other_inverts | 0.36 | 0.819 | 0.002 | 0.55 |
| AMT_mean ~ carrion_birds | 0.14 | 0.921 | 0.0002 | 0.56 |
| AMT_mean ~ piracy | -0.13 | 0.921 | 0.0002 | 0.56 |

### Supplementary discussion

#### Model limitations

While models such as ours are highly sensitive to parameter uncertainty, and may inadequately account for demographic species-specific variation (Dillingham & Fletcher, 2011; Richard et al., 2017), they remain a useful tool to guide conservation management actions (Niel & Lebreton, 2005; Robertson et al., 2014). There are a number of assumptions in the risk ratio model, including: the target species has constant adult survival, operates at low densities, $\lambda_{max}$ is constant across generations, and female fecundity is constant from age of first maturation (Dillingham, 2010; Dillingham et al., 2016; Niel & Lebreton, 2005). Further, our model is unable to capture species specific nuances in behaviour and life-stage, which will likely influence the resilience/risk of a species to a threat. For example, immature birds have a higher probability of dying in fisheries bycatch than breeding adults (Genovart et al., 2017). Similarly, young and immature birds are more likely to have higher loads of plastic ingested (van Franeker & Law, 2015). Some species are more gregarious when foraging, thus interactions with fisheries operations, or other human activities are likely to cause additional adult mortalities (Genovart et al., 2017). Abiotic factors also influence the level of risk at the individual level, for example, different levels of mortality are expected with the type of fisheries gear (Genovart et al., 2017). This means that our model may over- or underestimate the impact on a population, because the number of fatalities are attributed to breeding adults only and do not account for the variability in behaviour of individuals.

We assumed that changes in adult survival from marine threats was for breeding adults, thus do not account for non-breeding adults in the population (*i.e.*, adults on sabbatical, immature birds). Adult survival was assumed to be equal between sexes and monogamy was assumed (Genovart et al., 2017). We did not account for bias in sex and age classes, seasonality and cryptic mortality, which may underestimate the impact of marine threats on some populations (Mills & Ryan, 2005), although this was addressed to a degree for fisheries-related mortality in Richard and Abraham's (2013a) model.

In the same way that oceanic features vary across latitudes and water masses influencing resource distributions for seabirds, the intensity or existence of a threat is not distributed evenly (Ryan, 2016). Species that have large spatial distributions are likely to have variable population level responses to marine threats due to differences in spatial exposure, interspecific phenology, and dispersal patterns (e.g., climate change, 30). Complicating the strength of range as a risk predictor for seabirds is environmental stochasticity, which is closely linked to demographic stochasticity. That is the random variation of population dynamics due to discrete events (*i.e.*, changes to births and deaths from variable environmental factors, such as climate anomalies, prey availability (Tuljapurkar, 1990). Environmental stochasticity is widely recognised as being an important consideration in population growth models, particularly with small populations, where one event has the potential for catastrophic results (Weimerskirch, 2001). Thus, the effects of environmental stochasticity on vital rates for small populations, coupled with anthropogenic sources of adult mortality or reductions in reproductive output due to poor body condition (*i.e.*, plastic ingestion related) may be more pronounced (Lebreton & Clobert, 1991).

### Additional data table S1 (separate file)

**Extended data 1_colonyriskanalysis.csv**

Parameter estimates for the colony risk assessment for 37 colonies of 17 species and the impact of marine threats to recovery post-predator eradication.

### Additional data table S2 (separate file)

**Extended data 2_traitanalysis.csv**

Parameter estimates including 81 procellariform seabird species for the phylogenetic generalised least squares regression analysis to evaluate the influence of shared traits on the annual mortality threshold of a species.

**References**


Ashmole, N. P. (1971). Seabird ecology and the marine environment. *Avian Biology, 1*, 223–286.

Auman, H., Ludwig, J., Giesy, J., & Colborn, T. (1997). Plastic ingestion by Laysan albatross chicks on Sand Island, Midway Atoll, in 1994 and 1995. *Albatross Biology and Conservation*, 239–244. Retrieved from http://www.usc.edu/org/cosee-west/October06Resources/Related Articles/Plastic ingestion by Laysan Albatross chicks on Midway Atoll.pdf

Belda, E. J., & Sanchez, A. (2001). Seabird mortality on longline fisheries in the western Mediterranean: Factors affecting bycatch and proposed mitigating measures. *Biological Conservation*, *98*(3), 357–363.

Birdlife International. (2014). *Taxonomy*. Retrieved from http://www.birdlife.org/datazone/info/taxonomy

Birdlife International. (2016). *Species Distribution data*. Retrieved from http://datazone.birdlife.org/species/requestdis

Borrelle, S. B., Boersch-Supan, P. H., Gaskin, C. P., & Towns, D. R. (2016). Influences on recovery of seabirds on islands where invasive predators have been eradicated, with a focus on Procellariiformes. *Oryx*, *52*(2), 346–358.

Brooke, M. de L., Bonnaud, E., Dilley, B. J., Flint, E. N., Holmes, N. D., Jones, H. P., … Surman, C. (2018). Seabird population changes following mammal eradications on islands. *Animal Conservation*, *21*(1), 3–12.

Brooke, M. de L., O'Connell, T., Wingate, D., Madeiros, J., Hilton, G. M., & Ratcliffe, N. (2010). Potential for rat predation to cause decline of the globally threatened Henderson petrel Pterodroma atrata: Evidence from the field, stable isotopes and population modelling. *Endangered Species Research*, *11*(1), 47–59.

Caswell, H. (2001). *Matrix population models*. Wiley Online Library.

Caughley, G. (1994). Directions in conservation biology. *Journal of Animal Ecology*, 215–244.

Cornell Lab of Ornithology. (2019). *Birds of North America*. Retrieved from https://birdsna.org/Species-Account/bna/species/barpet/demography

Del Hoyo, J., Elliott, A., & Christie, D. (2011). Handbook of the Birds of the World. Vol. 15. Weavers to New World Warblers. *British Birds*, *104*, 225–228.

Dillingham, P. W. (2010). Generation time and the maximum growth rate for populations with age-specific fecundities and unknown juvenile survival. *Ecological Modelling*, *221*(6), 895–899. https://doi.org/10.1016/j.ecolmodel.2009.12.008

Dillingham, P. W., & Fletcher, D. (2011). Potential biological removal of albatrosses and petrels with minimal demographic information. *Biological Conservation*, *144*(6), 1885–1894. https://doi.org/10.1016/j.biocon.2011.04.014

Dillingham, P. W., Moore, J. E., Fletcher, D., Cortés, E., Alexandra Curtis, K., James, K. C., & Lewison, R. L. (2016). Improved estimation of intrinsic growth rmax for long-lived species: Integrating matrix models and allometry. *Ecological Applications*, *26*(1), 322–333. https://doi.org/10.1890/14-1990.1/suppinfo

Dunning, J. B. (2013). *Updates to the second edition of the CRC handbook of avian body masses: Https://ag. Purdue. Edu/fnr/Documents*. Retrieved from https://www.dropbox.com/sh/ykv8rino0lfziej/AABNHFRrKeODi_4qUtOVP2WLa?dl=0

Engen, S., & Sæther, B.-E. (2016). Optimal age of maturity in fluctuating environments under r- and K-selection. *Oikos*, *125*(11), 1577–1585. https://doi.org/10.1111/oik.03111

Frederiksen, M., Harris, M. P., Daunt, F., Rothery, P., & Wanless, S. (2004). Scale-dependent climate signals drive breeding phenology of three seabird species. *Global Change Biology*, *10*(7), 1214–1221.

Genovart, M., Doak, D. F., Igual, J., Sponza, S., Kralj, J., & Oro, D. (2017). Varying demographic impacts of different fisheries on three Mediterranean seabird species. *Global Change Biology*.

IUCN. (2017). *IUCN Red List of Threatened Species™* (Vol. 2017). Retrieved from http://www.iucnredlist.org

Jenouvrier, S. (2013). Impacts of climate change on avian populations. *Global Change Biology*, *19*(7), 2036–2057. https://doi.org/10.1111/gcb.12195

Jetz, W., Thomas, G., Joy, J., Hartmann, K., & Mooers, A. (2012). The global diversity of birds in space and time. *Nature*, *491*(7424), 444.

Laist, D. W. (1997). Impacts of marine debris: Entanglement of marine life in marine debris including a comprehensive list of species with entanglement and ingestion records. In *Marine Debris* (pp. 99–139). Springer.

Lebreton, J.-D., & Clobert, J. (1991). Bird population dynamics, management, and conservation: The role of mathematical modelling. In C. Perrins, J.-D. Lebreton, & G. Hirons (Eds.), *Bird population studies* (pp. 105–125). Retrieved from https://scholar.google.co.nz/scholar?q=Bird+population+dynamics%2C+management%2C++and+conservation%3A+the+role+of++mathematical+modelling&btnG=&hl=en&as_sdt=0%2C5

Mills, M. S. L., & Ryan, P. G. (2005). Modelling impacts of long-line fishing: What are the effects of pair-bond disruption and sex-biased mortality on albatross fecundity? *Animal Conservation*, *8*(4), 359-367. https://doi.org/10.1017/S1367943005002386

Morris, W. F., & Doak, D. F. (2002). Quantitative conservation biology. *Sinauer, Sunderland, Massachusetts, USA*.

Niel, C., & Lebreton, J.-D. (2005). Using demographic invariants to detect overharvested bird populations from incomplete data. *Conservation Biology*, *19*(3), 826–835. https://doi.org/10.1111/j.1523-1739.2005.00310.x

Oro, D. (2014). Seabirds and climate: Knowledge, pitfalls, and opportunities. *Frontiers in Ecology and Evolution*, *2*, 79.

Provencher, J., Borrelle, S. B., Sherley, R. B., Avery-Gomm, S., Hodum, P. J., Bond, A. L., … Mallory, M. L. (2018). Seabirds. In *World Seas, Volume III: Ecological Issues and Environmental Impacts.* Cambridge, MA, USA: Elsevier, Inc.

Richard, Y., & Abraham, E. R. (2013a). *Application of Potential Biological Removal methods to seabird populations* (Vol. 6480).

Richard, Y., & Abraham, E. R. (2013b). *Application of Potential Biological Removal methods to seabird populations* (Vol. 6480).

Richard, Y., & Abraham, E. R. (2013c). *Risk of commercial fisheries to New Zealand seabird populations* (Vol. 6480).

Richard, Y., Abraham, E. R., & Filippi, D. (2017). *Assessment of the risk of commercial fisheries to New Zealand seabirds, 2006-07 to 2014-15*. Ministry for Primary Industries, Manatū Ahu Matua.

Richard, Y., Abraham, E. R., & Filippi, D. P. (2011). Assessment of the risk to seabird populations from New Zealand commercial fisheries. *Final Research Report for Ministry of Fisheries Projects IPA 2009/19 and IP 2009/20*, 66.

Robertson, G., Moreno, C., Arata, J. A., Candy, S. G., Lawton, K., Valencia, J., … Suazo, C. G. (2014). Black-browed albatross numbers in Chile increase in response to reduced mortality in fisheries. *Biological Conservation, 169*, 319–333. https://doi.org/10.1016/j.biocon.2013.12.002

Rochman, C. M., Browne, M. A., Underwood, A. J., van Franeker, J. A., Thompson, R. C., Amaral-Zettler, L. A., … Amaral-Zettler, L. A. (2016). The ecological impacts of marine debris: Unraveling the demonstrated evidence from what is perceived. *Ecology, 97*(2), 302–312. https://doi.org/10.1890/07-1861.1

Rodríguez, A., Arcos, J. M., Bretagnolle, V., Dias, M. P., Holmes, N. D., Louzao, M., … Rodríguez, B. (2019). Future Directions in Conservation Research on Petrels and Shearwaters. *Frontiers in Marine Science*, *6*, 94.

Ryan, P. G. (1987). The effects of ingested plastics on seabirds: Correlation between plastic load and body condition. *Environmental Pollution*, *46*, 119–125.

Ryan, P. G. (2016). *Ingestion of plastics by marine organisms*.

Sæther, B.-E., & Engen, S. (2010). Population analyses. In A. P. Møller, W. Fiedler, & P. Berthold (Eds.), *Effects of climate change on birds*. Oxford, UK: Oxford University Press.

Salguero-Gómez, R., Jones, O. R., Archer, C. R., Bein, C., de Buhr, H., Farack, C., … Hoppe, G. (2016). COMADRE: a global data base of animal demography. *Journal of Animal Ecology*, *85*(2), 371–384.

Schreiber, E. A., & Burger, J. (2002). *Biology of Marine Birds*. CRC Press Boca Raton, Florida.

Tuljapurkar, S. (1990). *Population dynamics in variable environments*.

van Franeker, J. A., & Law, K. L. (2015). Seabirds, gyres and global trends in plastic pollution. *Environmental Pollution*, *203*, 89–96. https://doi.org/10.1016/j.envpol.2015.02.034

Vannela, R. (2012). Are We “Digging Our Own Grave” Under the Oceans? Biosphere-Level Effects and Global Policy Challenge from Plastic (s) in Oceans. *Environmental Science & Technology*, *46*(15), 7932–7933.

Weimerskirch, H. (2001). Seabird demography and its relationship with the marine environment. In E. A. Schreiber & J. Burger (Eds.), *Biology of Marine Birds (EA Schreiber and J. Burger, Eds.). CRC Press, Boca Raton, FL* (pp. 115–136).
